# Elucidating the Hierarchical Nature of Behavior with Masked Autoencoders

**DOI:** 10.1101/2024.08.06.606796

**Authors:** Lucas Stoffl, Andy Bonnetto, Stéphane d’Ascoli, Alexander Mathis

**Affiliations:** Ecole Polytechnique Fédérale de Lausanne (EPFL)

## Abstract

Natural behavior is hierarchical. Yet, there is a paucity of benchmarks addressing this aspect. Recognizing the scarcity of large-scale hierarchical behavioral benchmarks, we create a novel synthetic basketball playing benchmark (Shot7M2). Beyond synthetic data, we extend BABEL into a hierarchical action segmentation benchmark (hBABEL). Then, we develop a masked autoencoder framework (hBehaveMAE) to elucidate the hierarchical nature of motion capture data in an unsupervised fashion. We find that hBehaveMAE learns interpretable latents on Shot7M2 and hBABEL, where lower encoder levels show a superior ability to represent fine-grained movements, while higher encoder levels capture complex actions and activities. Additionally, we evaluate hBehaveMAE on MABe22, a representation learning benchmark with short and long-term behavioral states. hBehaveMAE achieves state-of-the-art performance without domain-specific feature extraction. Together, these components synergistically contribute towards unveiling the hierarchical organization of natural behavior. Models and benchmarks are available at https://github.com/amathislab/BehaveMAE.

## Introduction

Natural behavior is inherently hierarchical, i.e., it is built from spatially and temporally nested subroutines, as has been established in neuroscience and ethology (1–5). For example, a game of basketball is characterized by many part-whole relationships both across space (from groups to individuals to bodyparts and joints) and across time (from playing basketball to dribbling to taking a step). Here, we follow the nomenclature of Anderson and Perona (6), who delineate behavior into its elemental components: movemes, actions, and activities. Movemes represent the atomic units of behavior, e.g., a step and hence are akin to phonemes in language; actions comprise stereotypical combinations of movemes (e.g., dribbling, shooting, passing), while activities encompass species-typical sequences of actions and movemes, shaping adaptive or stereotyped behaviors (e.g., offensive plays, defending against fast breaks).

Skeletal action recognition and segmentation are common tasks in computer vision (7–14). However, they primarily focus on classifying actions at a single level of granularity, and state-of-the-art (SOTA) algorithms (14–16) recently saturated performance on popular (supervised) action recognition benchmarks (e.g., NTU and PKU-MMD (17–19)). Thus, there is a need for more challenging datasets including hierarchical action segmentation. To address this gap, we present two novel challenges: *Shot7M2*, a novel synthetic benchmark with annotated movemes, actions and activities for basketball play, and *hBABEL*, which extends BABEL (20) to hierarchical action segmentation.

Furthermore, we leverage Masked Autoencoders (MAEs) for modeling the hierarchical organization of behavior. MAEs have emerged as a powerful paradigm across many modalities (21–25). Building on this work, we propose a novel hierarchical model: hBehaveMAE, which fuses information across different spatio-temporal scales, enabling it to capture both fine-grained and coarse-grained features of behavior. We show that hBehaveMAE, contrary to the non-hierarchical variant, learns “interpretable latents” that decompose behavior into its hierarchical constituents (Fig. 1). There are many different meanings of interpretable in the literature (26, 27). By “interpretable latents” in hBehaveMAE, we refer to post-hoc human-interpretable explanations (26). Validating our design, we find a topographic mapping from architectural blocks to the behavioral hierarchy on Shot7M2 and hBABEL, i.e., lowest levels best explain movemes, while higher levels best explain actions and activities. Furthermore, hBehaveMAE reaches SOTA performance on the MABe22 animal benchmark (28).

**Fig. 1.**
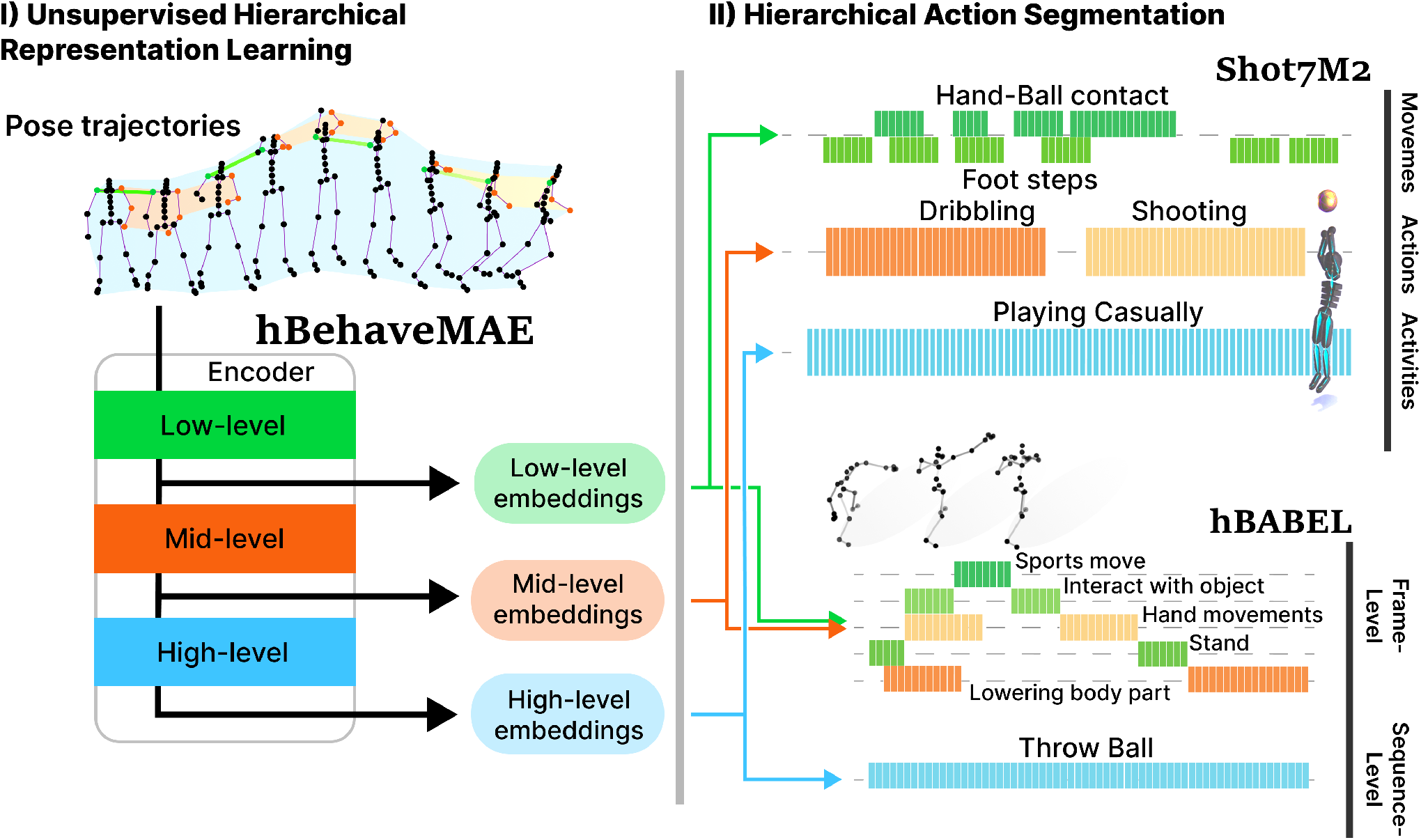
(I) Schematic of generalized, hierarchical Masked Autoencoder framework (hBehaveMAE) for hierarchical action segmentation that learns embeddings over several spatial and temporal scales. (II) We find that it captures aspects of the hierarchical nature of behavior, split into movemes (green), actions (orange) and activities (blue) on two novel benchmarks, Shot7M2 and hBABEL.

## Related Work

### Hierarchical Action Segmentation Benchmarks

Action recognition and segmentation benchmarks play a key role for developing algorithms to understand behavior. Existing benchmarks primarily focus on recorded human behaviors and often lack the necessary granularity to uncover the hierarchical nature of behavior (1–6). For instance, image-based datasets such as UCF (29), HMDB (30) and Kinetics400 (31) offer rich visual information, while skeleton-based datasets like NTU, PKU-MMD (18, 19), leverage pose data for better out-of-distribution generalization. However, these datasets primarily consist of exclusive actions and fail to capture compositional and hierarchical aspects of behavior. Recent benchmarks, such as MABe22 (28), addressed multi-animal behavior at two extreme timescales (e.g., actions and behavioral states such as day/night). Assembly101 (32) offers fine-grained and coarse-grained action segment annotations but is limited to hand poses and the supervised learning setting. Epic kitchens (33) comprises a wide range of action labels in a well-defined action segmentation benchmark but is only based on egocentric video data, while BABEL (20) annotates the AMASS motion capture dataset (34) with open-set behavioral annotations but primarily focuses on frame-level annotations for action recognition tasks, lacking hierarchical segmentation. Hence, there is a notable absence of hierarchical action segmentation benchmarks (Table 1) that we address.

**Table 1.** Comparison of skeletal action recognition and segmentation datasets with number of behaviors per frame and per individual. Duration per level is denoted in green, orange and blue (if available).

| Dataset | #Beh./Fr./Ind. | Behavior duration | Hierarchy | #Frames |
| --- | --- | --- | --- | --- |
| *Skeletal action recognition* |  |  |  |  |
| Human3.6M (35) | 1 | N.A. | ✗ | 3.6M |
| HumanAct12 (36) | 1 | $\approx 75$ | ✗ | 90.1k |
| NTU RGB+D-120 (19) | 1 | $\approx 70$ | ✗ | 8.0M |
| PKU-MMD (18) | 1 | 6 – 795 | ✗ | 5.4M |
| BABEL (20) | 1 | 150 | ✗ | 6.2M |
| *Skeletal action segmentation* |  |  |  |  |
| CalMS21 (37) | 1 | 2 – 1085 | ✗ | $\approx 2$M |
| MABe22 (mice) (28) | 2.43 | 1 – 1800 | ✗ | 4.7M |
| **Shot7M2** | 5.3 | 3 – 20 \| 12 – 400 \| 1800 | ✓ | 7.2M |
| **hBABEL** | 1.97 | 3 – 439 \| 128 – 1225 | ✓ | 6.2M |

Synthetic datasets have emerged as a promising avenue for creating large-scale, precisely annotated datasets for computer vision tasks (38–45). Building on the synthetic data paradigm, we introduce Shot7M2, a basketball playing dataset with an underlying hierarchical structure. Furthermore, we propose hBABEL, an extension of the BABEL dataset (20) tailored for hierarchical action segmentation. These benchmarks enable comprehensive evaluation of hBehaveMAE’s interpretability and performance across different hierarchical scales.

### Self-supervised Learning for Hierarchical Action Segmentation

Recent advancements in pose estimation and tracking have significantly advanced the study of behavior across various scientific domains (46–48). However, there remains a critical gap in both benchmarks and models for learning the hierarchical structure of behavior in an unsupervised fashion (49).

In applications, various computational approaches have been proposed to decompose behavior, described by pose trajectories, into “syllables” (50–56). However, these models typically operate at a single time-scale, which is an implicit or explicit parameter (56). In unsupervised representation learning competitions for behavioral analysis, such as MABe22 (28), adapted variants of BERT (57), Perceiver (58) and PointNet (59) reached strong results. Also, AmadeusGPT (60) performed well by generating task-program code from natural language user-input via language models. Bootstrap Across Multiple Scales (BAMS) (61) is the current SOTA method on MABe22 (28), and learns separate embedding spaces over two distinct time-scales. TS2Vec (62) incorporates hierarchy in the contrastive learning task, making these two approaches a valuable comparison for our hBehaveMAE models, which parse behavior at both temporal and spatial scales without relying on specialized feature extraction or training strategies.

### Hierarchical Masked Autoencoders

Originally introduced for Natural Language Processing (57), masked pretraining (63, 64) with transformers (65) has become the standard for self-supervised pre-training on large amounts of sequential, unlabeled data across fields, such as audio (66, 67), vision (21, 68, 69) or multi-modal signals (25, 70, 71). By only processing visible tokens during pre-training, Masked Autoencoders (MAEs) (21) significantly improved efficiency. These models have been expanded from spatial-only tasks, like reconstruction of images, to spatio-temporal data, such as videos (22, 23, 72) and skeletal pose data (15, 16, 73). Yan *et al*. (15) and Wu *et al*. (73) employ topological knowledge about human poses, and Mao *et al*. (16) leverage motion information to improve the masking strategy during pretraining. In contrast, due to the hierarchical architecture design, hBehaveMAE, learns hierarchical representations of behaviors.

Vanilla transformers struggle to encode hierarchical structures, as they primarily learn dependencies across different positions in sequential input data rather than capturing hierarchical relationships where the meaning of a token depends on its context within a broader structure (74). Efforts have been made to incorporate hierarchical structures into vision transformers (68, 75–79), with the idea of merging tokens at different levels in the architecture and utilizing local self-attention in lower levels. The recent Hiera model (80), which is based on hierarchical Vision Transformer (ViT) (81, 82), is designed for sparse MAE pretraining in a spatially hierarchical way and exhibits strong performance. In our work, we generalize this approach to the spatio-temporal domain with hBehaveMAE, emphasizing hierarchically interpretable representations for behavior.

### Hierarchical MAE for Behavior: h/BehaveMAE

We introduce h/BehaveMAE, a flexible MAE framework designed to capture the hierarchical structure of behavior from pose trajectories across various spatial and temporal scales (Fig 2). The proposed framework comprises two variants: the hierarchical model, referred to as *hBehaveMAE*, and its non-hierarchical counterpart, *BehaveMAE*. To collectively refer to both models, we use the notation *h/BehaveMAE*.

**Fig. 2.**
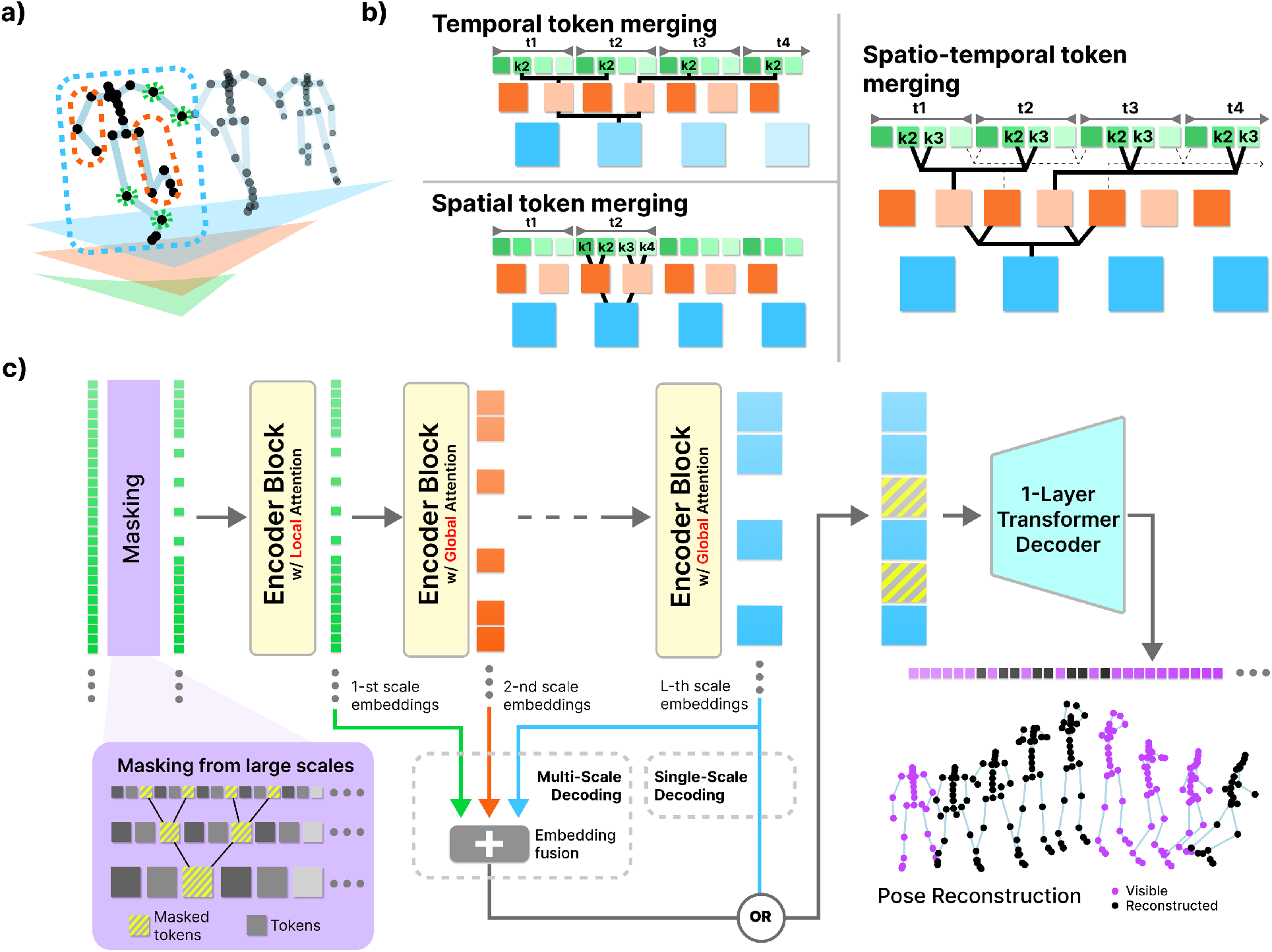
a) Input token representation. The input data consists of pose trajectories in 2D/3D from single or multiple individuals. This sequential data can be differently patched over spatial or temporal components, e.g., full pose over multiple frames (blue) or single keypoints over one timestep (green). These patches are encoded and flattened to form the input sequence to hBehaveMAE. **b) Hierarchical fusion**. Through the forward pass of hBehaveMAE models, the input tokens (green) can be fused either temporally, spatially or spatio-temporally, to form a pre-defined hierarchy. **c) Architecture of hBehaveMAE**. We mask a random set of tokens at the highest scale and back-project the mask to the lowest level to obtain the visible tokens in the input. At the end of every encoder block, that consists of multiple layers and uses either local or global attention, the tokens are combined according to a pre-defined fusion operation. A one-layer transformer decoder is taught to reconstruct the pose coordinates of the masked tokens, after receiving encoded visible tokens from the encoder, either combined over all blocks (multi-scale) or from the last layer only (single-scale).

#### MAE with Generalized Spatio-temporal Hierarchy

Let 𝒟 represent a set of unlabeled pose trajectories, each denoted by *γ* and comprising a sequence of poses 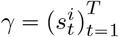. Here, each pose at time step *t* corresponds to the spatial coordinates of the *i*^*th*^ body part, typically represented in 2D 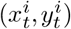 or 3D 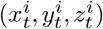. The length of each trajectory *T* is variable and often divided into sub-sequences during training, while the number of body parts *K* varies depending on the dataset. For instance, for 2D pose data, *γ* ∈ ℝ^*T*×*K*×*H*×*W*^, with frame height *H* and width *W*. For multi-individual scenarios, the *K*-dimension is expanded by stacking the postures of different individuals on top of each other, e.g., by augmenting the dimension to 3 · *K* for three individuals. We adapt spatio-temporal (video) MAEs (22, 23) to pose trajectories. The non-hierarchical version of our proposed model (**BehaveMAE**) is a natural extension of these, replacing input images with pose trajectories that are patched accordingly (Fig. 2a). Concretely, we first patch *γ* into space-time cubes, followed by embedding them via a learned linear projection. A random subset of tokens is masked and only the visible tokens are presented to the encoder. We use learned separable positional embeddings for the encoder, over time and space (e.g. different bodyparts) respectively. The final positional embedding added to the tokens is the sum of the two parts. A single-layer transformer decoder processes the encoded tokens alongside learnable mask tokens. The decoder’s primary computational task is to reconstruct the input sequence, with the loss computed solely over the masked tokens using a L2 loss (Supp. Mat. E.4 for loss comparison).

To learn hierarchical representations of behavior we introduce **Hierarchical BehaveMAE (hBehaveMAE)**, a spatio-temporal transformer architecture that adheres to the *vanilla* ViT style (83) while building on Hiera, a recent hierarchical model designed for images and videos (80). hBehaveMAE is tailored to accommodate spatio-temporal hierarchies and non-quadratic input data. While Hiera prioritizes efficiency and performance, our primary objective is the development of a generalized and interpretable model. Given our focus on learning a three-fold hierarchy across movemes, actions, and activities (6), a typical hBehaveMAE model comprises at least three blocks with a progressively increasing number of layers, hidden dimensions, and attention heads. Drawing from design principles for constructing functional hierarchies in images (80), our model incorporates local attention in the initial block. To facilitate hierarchical processing, we employ spatio-temporal fusion operators that dictate how embeddings are fused across time, individuals, and keypoints (Fig. 2b). Following each block, hBehaveMAE fuses tokens in either the temporal, spatial, or spatiotemporal dimension using query pooling attention (80). The temporal fusion operation can be expressed as:

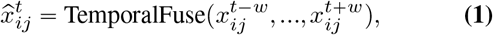

where 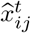 is the fused token at temporal position *t* with spatial position *ij*, and temporal fusion stride *w*. Similarly, the fusion operation in the spatial dimension is given by:

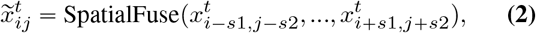

where 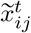 is the fused token at spatial position *ij* considering the spatial strides *s*1 and *s*2. The fusion operation in the spatio-temporal domain is defined as:

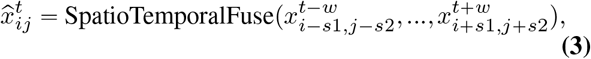

where 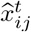 is the fused token considering both temporal and spatial relations.

Note that the spatial indices *i* and *j* may denote various entities such as a group of individuals, an individual pose, multiple keypoints constituting a body part, or a single keypoint, contingent upon the fusion’s hierarchical context. Thus, hBehaveMAE offers flexibility in both the construction of the initial hierarchy stage from pose trajectories (Fig. 2b), which may involve multiple individuals and the evolution of hierarchical fusion throughout its forward propagation.

#### Training and masking strategy

To facilitate self-supervised pre-training of h/BehaveMAE, we sample random patches without replacement from the set of embedded patches, akin to the masking strategy used in BERT (57) for 1D data, MAE (21) for 2D data, and the spatio-temporal MAE (22) for 3D data. The hierarchical nature of our model constrains the masking to be performed at the highest scale via a block masking scheme (84): the mask is projected to lower levels, in order to ensure consistent fusion throughout the forward pass of the model (Fig. 2c). Optimal performance for h/BehaveMAE is observed with masking ratios lower than the values commonly reported for video MAEs fine-tuned on coarse-grained downstream tasks such as action recognition (Fig. 6). Instead, h/BehaveMAE operates within the linear probing framework, decoding both fine-grained and coarse-grained features, thus necessitating lower masking ratios (see Ablations).

In summary, hBehaveMAE generalizes Hiera (80) to capture both temporal and spatial dependencies. In contrast to Hiera, that merges embeddings from all hierarchical stages to reconstruct pixels, we achieve strong performance with both single- and multi-scale decoding (Fig. 2c). Additionally, we simplified the decoder to a one-layer transformer to facilitate later testing of model embeddings through linear probing; we verified that stronger decoders did not improve (linear probing) performance (Supp. Mat. E.3).

### Hierarchical Action Segmentation Benchmarks

Filling the gap for hierarchical action segmentation we developed **Shot7M2** and **hBABEL**.

#### Shot7M2

We created **Shot7M2**, the **S**ynthetic, **H**ierarchical, and c**O**mpositional baske**T**ball dataset. Shot7M2 consists of 7.2 million frames designed to showcase hierarchical organization of basketball behavior (Fig. 3).

**Fig. 3.**
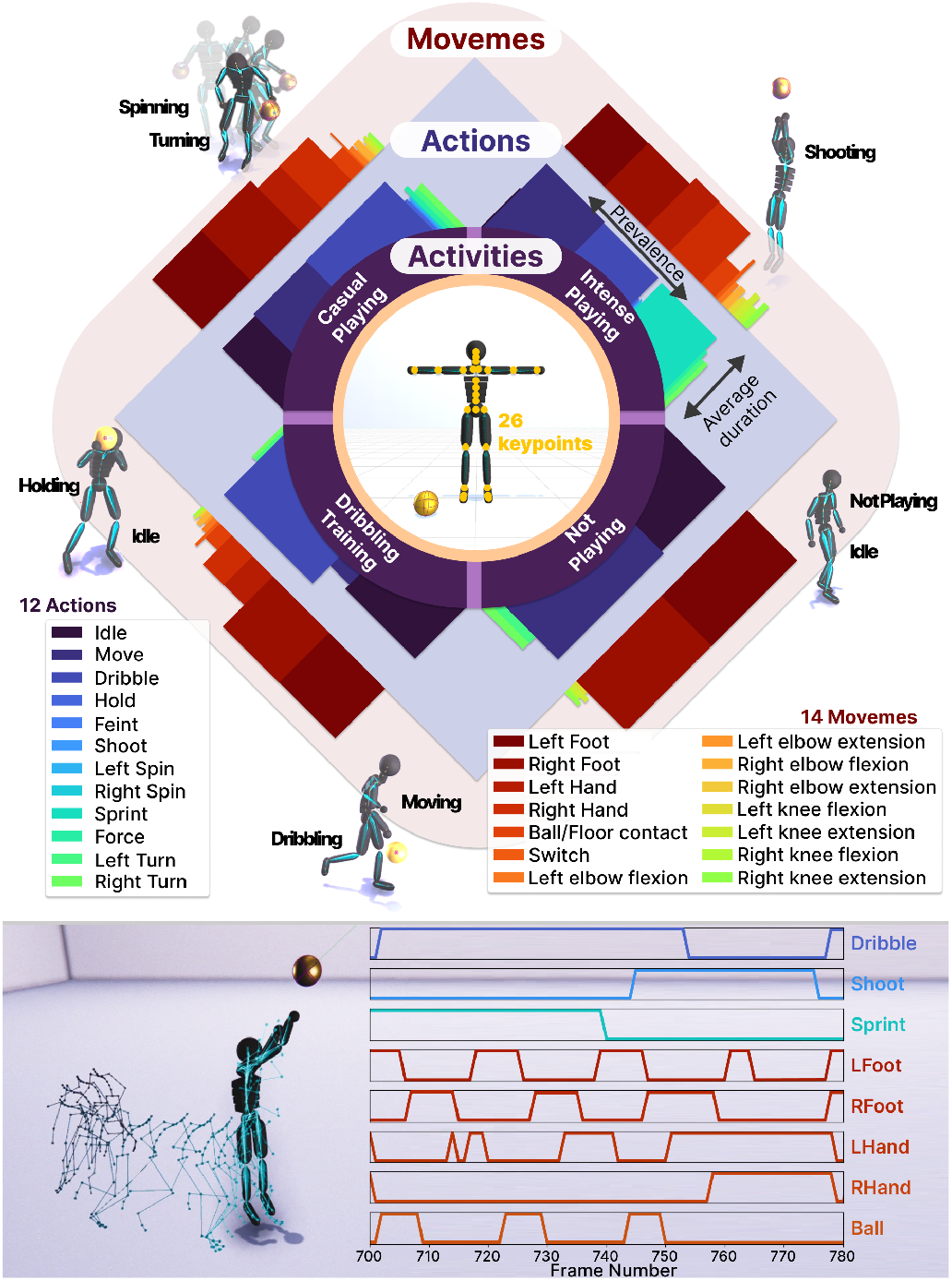
Overview of Shot7M2 dataset. The dataset contains 3D poses from 26 keypoints on a humanoid skeleton (center of the figure) and compositional behaviors from 4 activities with 12 actions and 14 movemes. Upper panel: statistics of the datasets: each activity constrains a specific prevalence and duration distribution for actions and movemes they are composed of (Supp. Mat. A.5). Histograms show this for each activity. Lower panel: example segment of one episode with (some of the) annotated behaviors.

#### Generation of Shot7M2

We generated 1000 2D trajectories per activity, sampled actions according to rules described thereafter and then based on the animation models by Starke *et al*. (85, 86) created 3D animations of a player following those paths and carrying out those actions. Action commands were randomly issued along the trajectory with activity-dependent frequencies. Movemes were extracted from the character’s kinematics and defined using different thresholds (Supp. Mat. A.3) for details on Shot7M2 generation).

#### Content of Shot7M2

Shot7M2 comprises 4000 episodes, each containing 1800 frames, where a single agent plays basketball.

Each episode is characterized by one of the four following activities: Casual play, Intense play, Dribbling training, Not playing. Each activity consists of actions from the following list: Idle, Move, Dribbling, Hold, Shoot, Feint, L/R Spin, Sprint, Force, L/R Turn. In addition, Shot7M2 contains movemes including hand-ball, ball-floor or foot-floor contact, flexion and extensions of the elbows and knees and switching the ball from one hand to the other. Shot7M2 exhibits hierarchical behavior representation across three levels: activities are defined for whole episode sequences, actions vary from 12 and 400 frames, and movemes range from 3 to 20 frames (Fig. 3). The compositionality of Shot7M2 is partly defined by its hierarchical nature, but also by the frequency of its behaviors. We also ensure a variable distribution of actions and movemes per activity by manipulating their prevalence and average duration across episodes (Supp. Mat. A.5 for statistics). By using Local Motion Phases (86), the animation generation is optimized to generate asynchronous movements which are translated into overlapping actions. In essence, Shot7M2 provides an opportunity to evaluate models designed for the analysis of human movement patterns and compositional actions.

#### hBABEL benchmark

We complement the synthetic dataset, by adapting BABEL as it has rich multi-level open-set behavioral annotations (20). BABEL provides sequence and frame-level behavioral annotations for the AMASS motion capture dataset (34), which contains over 45h of diverse human movements represented as SMPL-H mesh (87). For our hierarchical BABEL (**hBABEL**) benchmark, we predict the 3D skeletal poses from the vertices of the SMPL-H mesh and, as in the action recognition benchmark of BABEL (20), use the 25-joint skeleton format of NTU RGB+D (17). We further align each sequence according to their first frame using Procrustes alignment (88). The mean position across all sequence first frames is used as a reference frame for alignment.

hBABEL has a hierarchical nature since it is described by frame-level labels, which are components of sequence-level labels (20). We processed the behavioral annotations in the same manner as in prior work (89–91). To ensure a similar distribution of labels in the training and testing set during evaluation, we counted the number of segments and selected the top 120 most frequent behaviors for the frame-level subtasks and the top 60 most frequent actions for the sequence-level subtasks. Note that this implies that some episodes do not have any annotations.

Details can be found in the supplementary materials (Section A).

### Experiments

We performed experiments on three different datasets: the 3-level synthetic dataset Shot7M2, the hierarchical action segmentation variant of BABEL (20), hBABEL, and the 2022 Multi-Agent behavior Challenge (MABe22)

#### Benchmarking Datasets and Implementation Details

**MABe22** contains a collection of mouse triplet video clips selected for the analysis of representation learning algorithms. It consists of 5336 60-second clips capturing three mice at a rate of 30Hz. Each clip includes trajectory data that represents the postures and movements of the mice, which is obtained by tracking a set of 12 anatomically defined keypoints in 2D. The dataset contains 13 actions, that are annotated either at the frame level or the sequence level. The labels encompass various aspects including human annotations and experimental setups and include biological variables (e.g., animal strain) environmental factors (e.g., time of day), or social behaviors (e.g., chasing, huddling).

**Shot7M2** contains 4000 sequences of 1800 frames for a total of 7.2M frames. The skeleton of the individual is defined at all time and consists of 26 keypoints in 3D. Including 4 activities, 12 actions and 14 movemes, Shot7M2 describes 30 densely annotated non-exclusive behaviors. Following the protocol for MABe22 (28), 32% of the dataset is used for pre-training, while 68% is used for evaluation. The dataset was randomly separated by episode while ensuring a good partition of activities.

**hBABEL** extends BABEL (20) that provides textual descriptions for the motion sequences in the AMASS collection (34). Following the official train, val, and test splits, the dataset comprises 6601, 2189 and 2079 sequences, respectively. We filtered out sequences shorter than 0.5 seconds (to keep long enough sequences for the encoder) and use the updated text annotations from TEACH (90). For both the frame and sequence annotations, we make use of the categorical action labels. Human motions in hBABEL are described by 25 3D joint coordinates, following the NTU RGB+D format (17).

#### Training protocols

The pre-training configuration of h/BehaveMAE is as follows. We train for 200 epochs (including 40 warmup epochs) using the AdamW optimizer (92) with a learning rate of 1.6 ^− 4^ (Supp. Mat. B). For all our experiments, we use random masking (for hBehaveMAE based on the highest scale). For MABe22, we follow the data augmentation scheme of (93) and use reflections, rotations and Gaussian noise added to the keypoints. For Shot7M2 and hBABEL, we do not employ any data augmentation (for any model). The joint positions are normalized by the size of the grid for MABe22, and projected to egocentric coordinates for hBABEL and Shot7M2.

#### Baselines

We implemented five, diverse baselines on Shot7M2 and hBABEL. To simply account for frequency, we evaluated a *base* classifier, which always predicts the most probable outcome. We trained a Principal Component Analysis (PCA) model on each individual frame along with a temporal version of PCA, which includes all information from a temporal window of 30 or 5, for Shot7M2 and hBABEL, respectively (PCA-30 and PCA-5). We trained a Trajectory Variational AutoEncoder (TVAE) (94) model on 300 epochs using a learning rate of 1*e* ^− 5^ and a temporal stride of 30. To incorporate SOTA methods from MABe22 into Shot7M2 and hBABEL, we trained TS2Vec (62) & BAMS (61) without making use of additional features to make it comparable to h/BehaveMAE (Supp. Mat. B).

#### Evaluation

The evaluation on all datasets is based on linear probing and follows the evaluation protocol of the MABe22 challenge (28). Considering an evaluation set, which is independent from the data used in pre-training, a linear classifier is trained on top of the frozen representations, independently for each behavior on 75% of the evaluation set and evaluated on the remaining 25%. For hBABEL, we group the scores by averaging over the top 10, 30, 60 and 90 most frequent behaviors for frame-level subtasks and over the top 10, 30 and 60 for the sequence-level tasks. For MABe22 mice the maximum size of a frame embedding is given by the challenge, i.e. 128, while for Shot7M2 and hBABEL, we decided to allow a size of 64; increasing the embedding size improves performance (Supp. Mat. E.5).

When we have multiple embeddings per frame (e.g. multiple mice in MABe22 or multiple bodyparts in Shot7M2 or hBABEL) we perform average pooling and, if needed, compress the embedding with PCA.

### Comparison to State-of-the-Art

#### MABe22

First, we evaluated h/BehaveMAE on the mouse task of MABe22 (28). hBehaveMAE achieves SOTA results and outperforms previous methods in both sequence-level and frame-level classification tasks (Table 2), with similar performance to SOTA on regression tasks. While these algorithms use additional contrastive learning objectives (T-PointNet, T-BERT), additional supervision (T-Perceiver and T-BERT) or additional behavioral features (T-Perceiver, T-PointNet, T-BERT, BAMS), h/BehaveMAE is trained exclusively on the raw pose trajectories. We will also evaluate TS2Vec (62) and BAMS (61) on Shot7M2 and hBABEL, as both models perform well and incorporate temporal hierarchy.

**Table 2.** Comparison to SOTA methods on MABe22 Mice Triplets. Models are split according to their pre-training set: training set only (inductive) or all available data (transductive).

| Method | feat. | All MSE (↓) | All F1 | Frame F1 | Seq. F1 |
| --- | --- | --- | --- | --- | --- |
| *Inductive setting* |  |  |  |  |  |
| Base |  | .0941 | 21.5 | 13.2 | 44.4 |
| PCA |  | .0943 | 19.6 | 12.1 | 53.1 |
| TVAE (28, 94) |  | .0942 | 22.7 | 15.5 | 54.9 |
| TS2Vec (62) |  | .0940 | 24.5 | 16.3 | 61.4 |
| T-Perceiver (28, 58) | × | .0933 | 27.1 | 17.6 | 69.7 |
| T-PointNet (28, 59) | × | .0930 | 28.2 | 19.6 | 66.6 |
| T-BERT (28, 57) | × | .0927 | 29.7 | 19.9 | 73.7 |
| BAMS (61) | × | **.0912** | 29.9 | 19.5 | 76.9 |
| BehaveMAE |  | <u>.0913</u> | <u>30.5</u> | **20.1** | <u>77.4</u> |
| hBehaveMAE |  | .0918 | **30.7** | **20.1** | **78.5** |
| *Transductive setting* |  |  |  |  |  |
| T-GPT (28, 95) |  | .0933 | 27.4 | 19.1 | 64.9 |
| BAMS (61) | × | **.0904** | 30.8 | 19.8 | 80.1 |
| BehaveMAE |  | .0906 | 32.4 | 21.5 | **81.0** |
| hBehaveMAE |  | .0918 | **32.9** | **22.3** | <u>80.3</u> |

#### Shot7M2

hBehaveMAE shows the best performances for activities, actions and movemes on Shot7M2 with averaged F1-scores of 80.9%, 58.5% and 67.3%, respectively. Importantly, it shows stronger performances than the non-hierarchical architecture BehaveMAE (Table 3). We also found that PCA was surprisingly competitive.

**Table 3.** Results on Shot7M2: hBehaveMAE outperforms baseline methods on all three action categories, with large gains on activities and actions. The *All F1* score is calculated as the average over each behavioral scale average score. BehaveMAE scores are obtained from embeddings of layer 4 (its best). hBehaveMAE scores are obtained from the maximum over its embedding layers.

| Method | All F1 (↑) | Movemes | Actions | Activities |
| --- | --- | --- | --- | --- |
| Base | 21.8 | 18.0 | 18.6 | 40.0 |
| PCA | 60.8 | 61.8 | 51.0 | 69.5 |
| PCA-30 | 61.9 | 66.7 | 50.8 | 68.3 |
| TVAE (28, 94) | 51.2 | 41.0 | 47.0 | 65.5 |
| TS2Vec (62) | 59.6 | 60.2 | 50.0 | 68.6 |
| BAMS (61) | 53.7 | 48.6 | 45.3 | 67.1 |
| **BehaveMAE** | **64.3** | **61.0** | **55.1** | **76.7** |
| **hBehaveMAE** | **64.7** | **67.3** | **58.5** | **80.9** |

#### hBABEL

BehaveMAE performs best both on the frame-level subtasks with an average Top 30 F1-score of 20.3% and on the sequence-level subtasks with an average Top 10 F1-score of 23.4 (Table 4). Even though BAMS showed strong performances on the MABe22 benchmark, we encountered difficulties to optimize it on hBABEL (Supp. Mat. B.3). Again, PCA was surprisingly competitive.

**Table 4.**
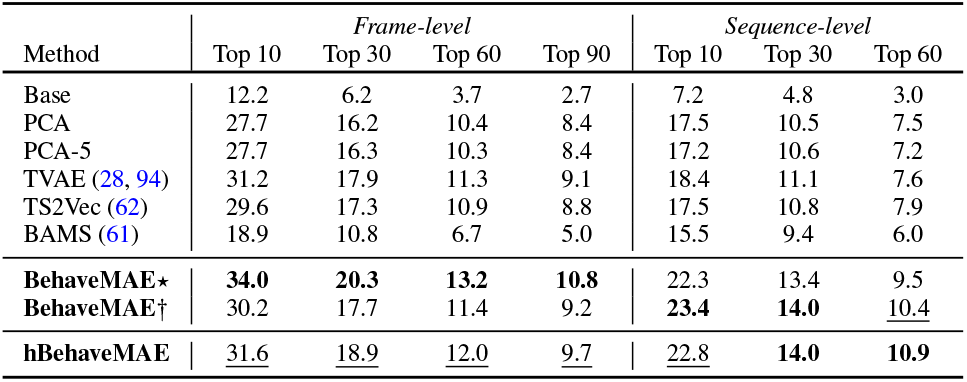
Results on hBABEL: BehaveMAE excels in either frame-level or sequence-level tasks, depending on the model setting, while hBehaveMAE effectively balances performance across both scales. ⋆denotes 15×1×15 token input for BehaveMAE (grouped bodyparts over 15 frames), while † indicates 15×1×75 (full pose over 15 frames). BehaveMAE scores are obtained from embeddings of layer 5 (its best).

| Method | Frame-level |  |  |  | Sequence-level |  |  |
| --- | --- | --- | --- | --- | --- | --- | --- |
|  | Top 10 | Top 30 | Top 60 | Top 90 | Top 10 | Top 30 | Top 60 |
| Base | 12.2 | 6.2 | 3.7 | 2.7 | 7.2 | 4.8 | 3.0 |
| PCA | 27.7 | 16.2 | 10.4 | 8.4 | 17.5 | 10.5 | 7.5 |
| PCA-5 | 27.7 | 16.3 | 10.3 | 8.4 | 17.2 | 10.6 | 7.2 |
| TVAE (28, 94) | 31.2 | 17.9 | 11.3 | 9.1 | 18.4 | 11.1 | 7.6 |
| TS2Vec (62) | 29.6 | 17.3 | 10.9 | 8.8 | 17.5 | 10.8 | 7.9 |
| BAMS (61) | 18.9 | 10.8 | 6.7 | 5.0 | 15.5 | 9.4 | 6.0 |
| **BehaveMAE\*** | **34.0** | **20.3** | **13.2** | **10.8** | 22.3 | 13.4 | 9.5 |
| **BehaveMAE†** | 30.2 | 17.7 | 11.4 | 9.2 | **23.4** | **14.0** | **10.4** |
| **hBehaveMAE** | **31.6** | **18.9** | **12.0** | **9.7** | **22.8** | **14.0** | **10.9** |

We emphasize that hBABEL is challenging due to the high number of behaviors, the sparsity of these behaviors over the evaluation dataset and the high variation of behavior durations.

We note that SOTA methods on the BABEL action recognition benchmark (20) achieved only an F1 score of 41.1% (24.5% normalized by frequency) despite the fact that supervised action recognition of short clips is an easier task than hBABEL’s proposed action segmentation through linear probing. On the positive side, this opens up ample research possibilities, e.g., with language models (60).

#### Learning the Hierarchy of Behavior

Next we delve into the hierarchical organization of behavior as learned by hBehaveMAE. First, we present a comparative analysis, revealing the interpretability of hBehaveMAE’s hierarchical architecture compared to the non-hierarchical counterpart with the same number of overall layers (9 - which are distributed over 3 blocks in hBehaveMAE). On both Shot7M2 and hBABEL, scores for low-level actions (movemes and Per-Frame actions) are better decoded from the the first level embeddings of hBehaveMAE, while higher level actions (actions, activities or sequence actions) are better decoded from the higher-level embeddings of hBehaveMAE (Fig. 4). This shows that hBehaveMAE is able to effectively learn a hierarchical structure of the action categories, which validates that the fusion operation is essential. Second, we carry out a similar analysis over a wide range of block sizes (Fig. 5). We observe that early layers of the model are best suited for decoding movemes, while late layers excel in capturing activities, independent of the number of hierarchical blocks; we also found this without linear probing when performing a clustering analysis (Supp. Mat. D).

**Fig. 4.**
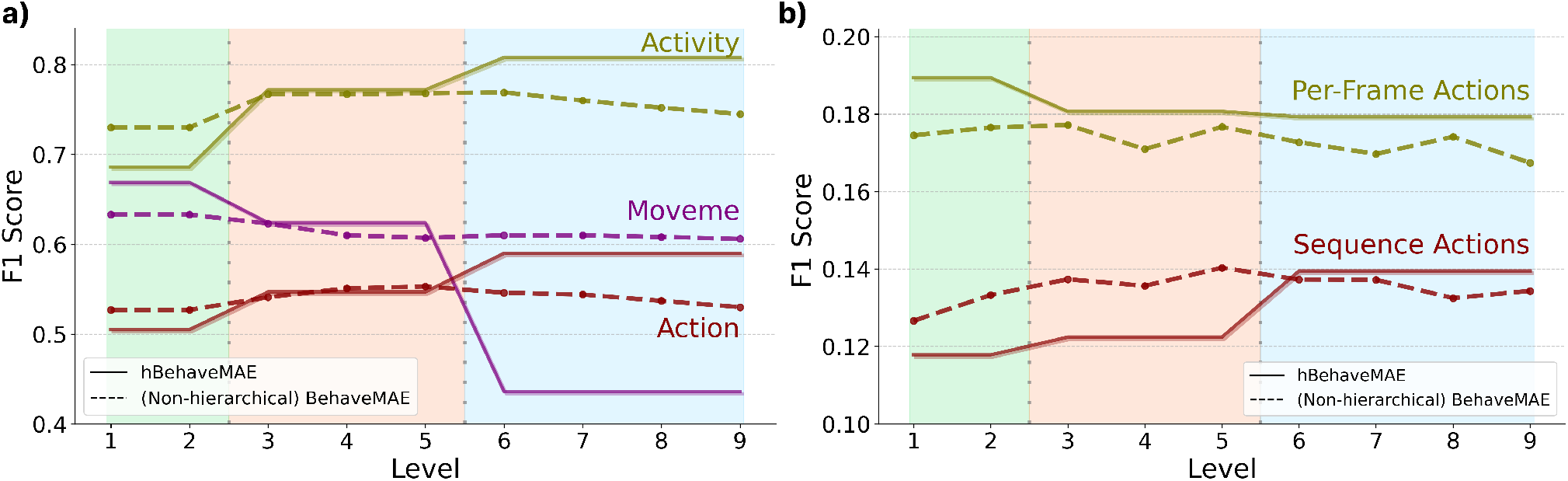
Performance and interpretability of hBehaveMAE (solid) vs. BehaveMAE (dashed) per layer. a) Shot7M2. hBehaveMAE outperforms BehaveMAE on all three groups (movemes, actions, activities) while showing “interpretable” performance compared to the relatively flat curves of BehaveMAE. **b) hBABEL (top 30 F1)**. hBehaveMAE better balances overall performance with lower layers better decoding frame-level actions and higher blocks sequence-level actions. Background colors indicate hierarchical blocks.

**Fig. 5.**
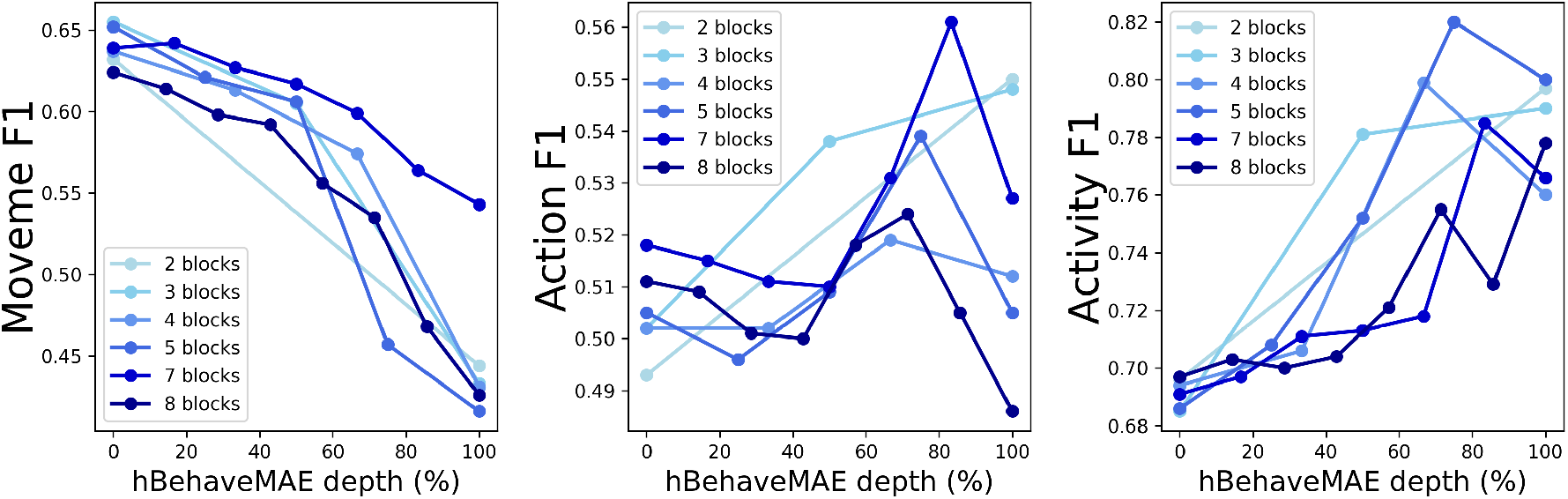
Impact of depth on hBehaveMAE interpretability highlighting early layers are robustly best for movemes and late layers best for activities. Models are tested on Shot7M2 and the overall hierarchical stride is kept the same (8×1×24), independent of the number of blocks.

#### Ablations

We test key design choices of h/BehaveMAE using the Shot7M2 benchmark and linear probing against movemes, actions, and activities.

#### Masking ratio

Following insights from Ryali *et al*. (80), who observed that hierarchical MAEs for vision benefit from slightly lower masking ratios compared to standard MAEs (due to the increased difficulty of the pretext task coming from the masking at highest scale), we investigated the effect of masking ratio variation on h/BehaveMAE (Fig. 6). Lower masking ratios enhance the model’s ability to decode fine-grained movements, resulting in higher performance on movemes and actions. Conversely, higher masking ratios are advantageous for capturing activity-level patterns, which aligns with observations from video MAEs (22, 23), which prioritize learning latents for sequence classification tasks. We determined the optimal masking ratio across scales on Shot7M2 to be around 70%.

**Fig. 6.**
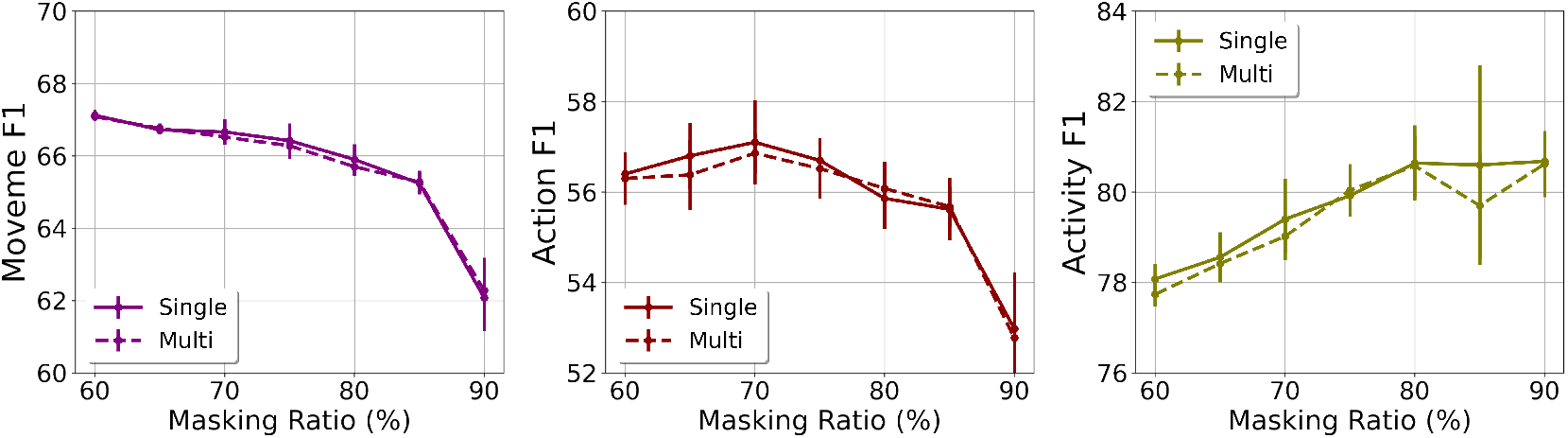
Masking ratio and single-scale decoding. The optimal masking ratio is obtained around 70% with effectively balancing performance on movemes and actions (60-75%) and activities (70-90%), while using single-scale information (from the last layer) during training performs on par with multi-scale decoding. Experiments were conducted with 5 different random seeds for robustness.

#### Singe-scale decoding

Experimenting with single-scale and multi-scale decoding (Fig. 2c), we find that both decoding strategies perform on par, which is in contrast to Hiera (80) (Fig. 6).

#### Inductive bias of attention

Restricting the model to utilize only local attention resulted in performance drops across all categories, particularly on actions and activities. Conversely, exclusively using global attention, leading to higher computational costs, decreased performance on actions and activities, likely because of the over-reliance on fine-grained information, which impedes the learning of effective abstractions. The best performance was achieved by employing local attention in the lower blocks (Table 5).

**Table 5.**
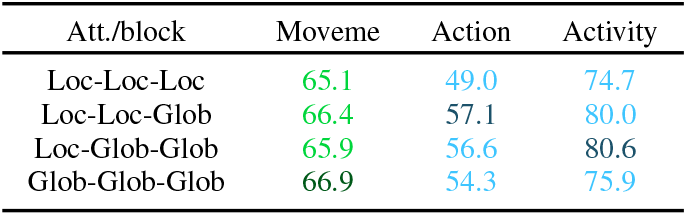
Local attention. hBehaveMAE benefits from local attention in its lower blocks. F1 scores are obtained from best block (green: 1st; blue: 3rd).

Additional ablations (reconstruction target, loss function, encoder and decoder sizes and the droppath rate) show the robustness of hBehaveMAE (Supp. Mat. E).

## Conclusions and Limitations

We make two key contributions. Firstly, we introduce the first hierarchical action segmentation benchmarks: Shot7M2 and hBABEL. Due to its synthetic nature, Shot7M2 might contain unnatural movements and hBABEL only has two annotated levels. Secondly, we developed h/BehaveMAE, a framework for discovering behavioral states from raw pose data, whose performance on these challenging benchmarks can still be improved. While the structural hierarchy in our model’s architecture needs to be pre-defined, the functional hierarchy was found to emerge naturally and robustly from the data (Fig. 5). We hope that our work motivates others to create hierarchical action segmentation benchmarks and models.

## Acknowledgements

We thank Haozhe Qi, Alberto Chiappa, Alessandro Marin Vargas, Adriana Perez Rotondo, and Niels Poulsen for feed-back. We acknowledge funding from EPFL’s AI4science program (SA, AM), the Microsoft Swiss Joint Research Center and EPFL.

## Supplementary Materials

### Supplementary Note A: Details on Hierarchical Action Segmentation Benchmarks

#### A.1. What is hierarchical behavior?

Certain (non-exclusive) conditions are desired for a dataset to be called hierarchical:

1. *Multiple behavior instances*: The dataset should contain instances where multiple behaviors occur simultaneously within a single frame or sequence (e.g., walking and talking).
2. *Nested behaviors*: The dataset should exhibit nested behaviors, where larger behaviors (e.g., activities) contain smaller, more granular behaviors (e.g., actions). For instance, the activity of “cleaning” will typically involve many different actions. In basketball, “attack” will comprise many potential actions, but note that there is no necessary action. One can even participate in an attack, without dribbling, touching the ball or running.
3. *Temporal dependencies*: Behaviors should exhibit temporal dependencies, where the occurrence of one behavior influences the likelihood or timing of another behavior.
4. *Varied duration distributions*: Behaviors should have varying duration, allowing for actions to be composed of multiple sub-actions or movements.

#### A.2. Dataset comparisons

In the main text, we emphasized the lack of existing hierarchical skeletal-based action segmentation datasets. To study hierarchical behavior, a dataset with densely annotated labels (with more than one behavior per frame and per individual) is essential, encompassing diverse duration distributions to accommodate, e.g., actions within activities or movemes within actions (following the nomenclature of Anderson and Perona (6)). For scalability reasons, we are particularly interested in skeletal input data and thus compiled a comparative table showcasing publicly available skeletal action recognition and action segmentation datasets (Table 1 in the main text). The table highlights that existing action segmentation benchmarks lack annotations of multiple behaviors at the same time (conjunctive). Yet, one could be sitting and writing or standing and writing. Our novel benchmarks hBABEL and especially Shot7M2 address this compositionality and hierarchy.

#### A.3. Generation details for Shot7M2

Grounded from the animation models by Starke et al. (85, 86), we use a Neural State Machine composed of a Motion Prediction Network and a Gating Network to predict future character poses from the current pose, set of control commands, goal position, environment geometry and a set of actions. The Gating Network modulates the weights of the Motion Prediction Network using a mixture of experts to ensure smooth transitions between actions. Additionally, low level controls normally given by a gamepad input are auto-encoded through a Generative Control Scheme to produce more refined control commands.

For each activity, except Dribbling Training, we generated 1000 2D-trajectories each in a constrained environment using down-sampled Brownian motion. Each trajectory is complemented with a set of control commands based on the motion direction and the average speed of the character. The ball control commands alternate between front, left, right and circular motion in both directions with respect to the character’s root orientation. Actions responsive to commands such as Hold, Left Spin, Right Spin or Shoot are randomly called along the trajectory. In Dribbling Training, all trajectories follow either a linear trajectory with various amplitude of sinusoidal lateral motions, or a static position where the player only dribbles at the same location. There are no Hold or Shoot commands in the Dribbling Training activity. During the Intense Playing activity, the Sprint action is maintained during most of the episode with some breaks lasting from 40 to 80 frames. In the Not Playing activity, the character never carries the ball and only follows a trajectory. In order to grasp the ball after a shooting action, the player uses a special power, defined by an action named Force, to attract the ball toward themselves. Thus, the trajectories together with sampled action sequences are turned into 3D animations of a player following those paths and carrying out those actions via Starke’s models (85, 86).

The code to generate and the link to download Shot7M2 are available at https://github.com/amathislab/BehaveMAE.

**Fig. S1.**
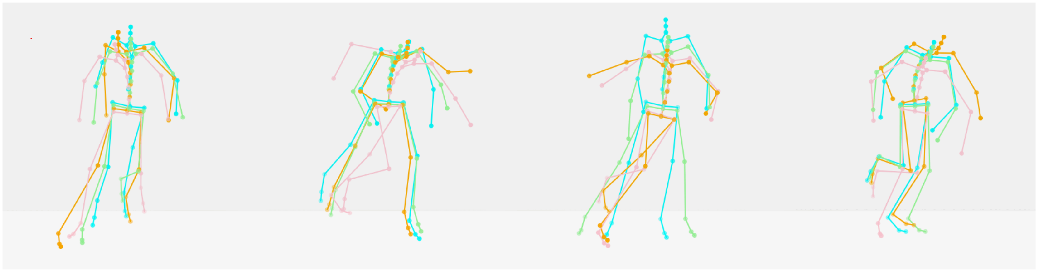
Variability of Shot7M2 poses. Four example frames taken from the sequence of actions and trajectories but with different pink noise seeds (colors).

#### A.4. Increasing the variability of 3D poses in Shot7M2

We motivate the need for adding noise into the pose by the deterministic nature of the trained Neural State Machine (86), which tends to create a linear mapping between the pose information and the combination of behaviors. To introduce variability while maintaining naturalistic poses for the same combination of actions, we employed pink noise in the input of the Neural State Machine model. We picked pink noise, characterized by time-correlated white noise, to prevent temporal jittering. Noise modulation was applied to all keypoints except the feet to ensure precise contact with the ground, which is defined as a moveme. The noise values for each timestep were generated at the start of each episode and influenced the original pose without disrupting action execution (Figure S1).

#### A.5. Statistics on Shot7M2 behavior labels

Here we provide additional details on the Shot7M2 benchmark (Table S1). Shot7M2 exhibits a high density of actions, averaging 5.3 behaviors per frame (Table 1). The dataset varies in scale between the distribution of durations of actions and Movemes (Figure S2), with the duration of activities fixed at 1800 frames. Additionally, the distribution of actions and Movemes varies depending on the player’s activity (Figure S3 and S4), and the number of Movemes varies depending on the player’s Action (Figure S5). These aspects underscore the hierarchical nature of the dataset.

**Fig. S2.**
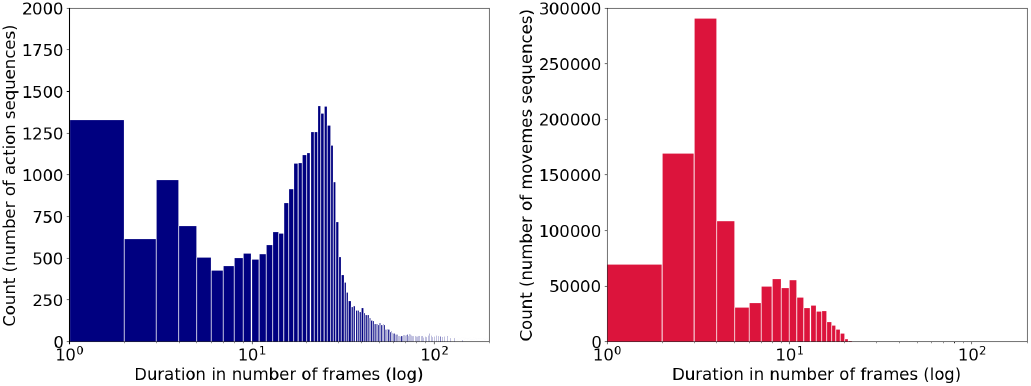
Number of sequences per sequence length for all actions (left) and all movemes (right).

**Fig. S3.**
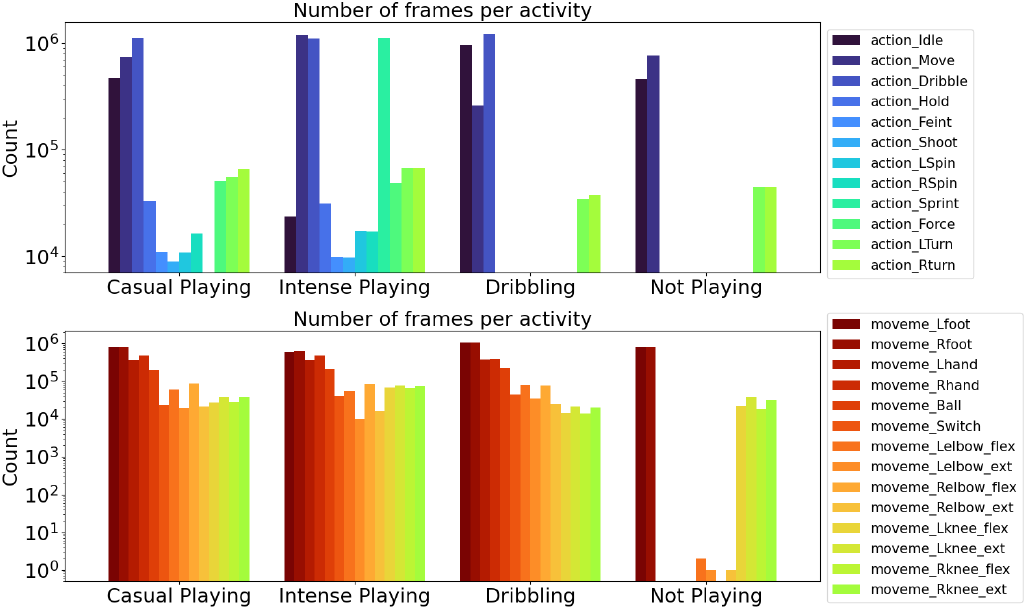
Number of frames per action (top) and movemes (bottom) for all 4 activities.

**Fig. S4.**
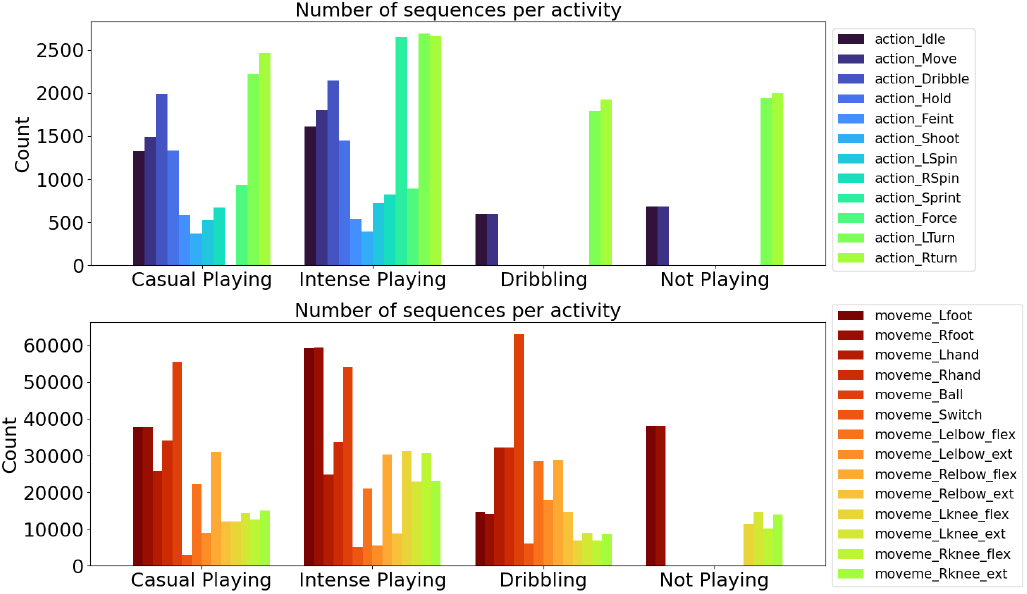
Number of sequences per action (top) and movemes (bottom) for all 4 activities.

**Fig. S5.**
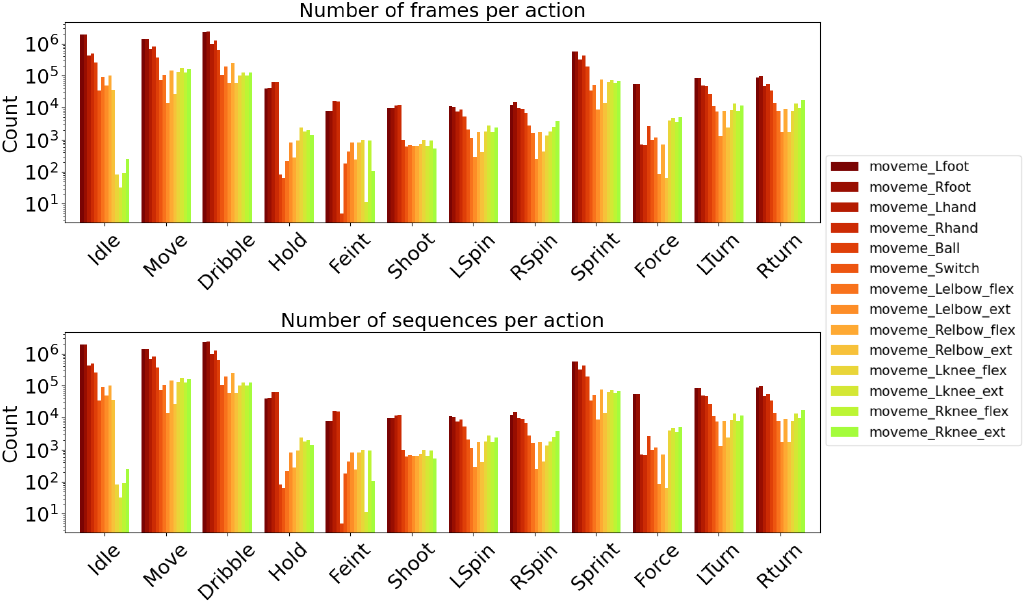
Number of frames (top) and number of sequences (bottom) per moveme for all 12 actions.

**Table S1.** Duration and prevalence of behaviors in Shot7M2.

| Behaviors | Duration (frames) | Number of frames |
| --- | --- | --- |
| *Activities* |  |  |
| Casual | 1800.0 ± 0.0 | 1,800,000 |
| Intense | 1800.0 ± 0.0 | 1,800,000 |
| Dribbling | 1800.0 ± 0.0 | 1,800,000 |
| Not Playing | 1800.0 ± 0.0 | 1,800,000 |
| *Actions* |  |  |
| Idle | 669.59 ± 1,261.29 | 2,816,275 |
| Move | 954.32 ± 879.31 | 4,361,254 |
| Dribble | 1230.07 ± 28,029.71 | 5,082,671 |
| Hold | 34.03 ± 50.32 | 94,578 |
| Feint | 27.38 ± 8.82 | 30,660 |
| Shoot | 36.01 ± 12.9 | 27,340 |
| LSpin | 33.06 ± 60.25 | 41,128 |
| RSpin | 32.75 ± 55.06 | 49,030 |
| Sprint | 626.47 ± 720.99 | 1,658,876 |
| Force | 80.34 ± 91.19 | 146,462 |
| LTurn | 34.26 ± 28.38 | 296,053 |
| RTurn | 35.03 ± 28.41 | 316,672 |
| *Movemes* |  |  |
| Left foot | 32.09 ± 182.65 | 4,801,318 |
| Right foot | 32.4 ± 184.6 | 4,840,360 |
| Left hand | 19.47 ± 11.49 | 1,616,040 |
| Right hand | 19.66 ± 11.44 | 1,969,050 |
| Ball | 5.37 ± 1.59 | 927,054 |
| Switch | 11.15 ± 5.03 | 158,731 |
| Left elbow flexion | 4.03 ± 0.85 | 289,528 |
| Left elbow extension | 2.91 ± 1.07 | 94,393 |
| Right elbow flexion | 3.99 ± 0.88 | 358,224 |
| Right elbow extension | 2.63 ± 0.87 | 94,147 |
| Left knee flexion | 3.15 ± 1.24 | 194,069 |
| Left knee extension | 4.16 ± 1.26 | 253,619 |
| Right knee flexion | 3.1 ± 1.22 | 187,660 |
| Right knee extension | 3.99 ± 1.31 | 241,444 |

#### A.6. Calculation of F1 scores in Shot7M2

The F1 scores have been calculated independently for each activity, action and moveme. For clarity, we averaged the F1 scores over the following pairs before reporting them (e.g., in Table S9):

- Left turn + Right turn → Turn
- Left spin + Right spin → Spin
- Left knee extension + Left knee flexion → Left knee
- Right knee extension + fight knee flexion → Right knee
- Left elbow extension + Left elbow flexion → Left elbow
- Right elbow extension + Right elbow flexion → Right elbow

#### A.7. Statistics on hBABEL behavior labels

We believe that the action segmentation benchmark hBABEL also demonstrates a hierarchical nature albeit to a lesser degree than Shot7M2. It features non-exclusive sets of actions with a density of 1.97. Moreover, it encompasses two distinct scales of behavior duration (Figure S6) and varying distributions of frame-level actions per sequence-level actions (Figure S7). We note that frame-level and sequence-level actions do not map (in a clear cut way) onto movemes, actions and activities. Thus, we use this distinct nomenclature from Babel (20).

**Fig. S6.**
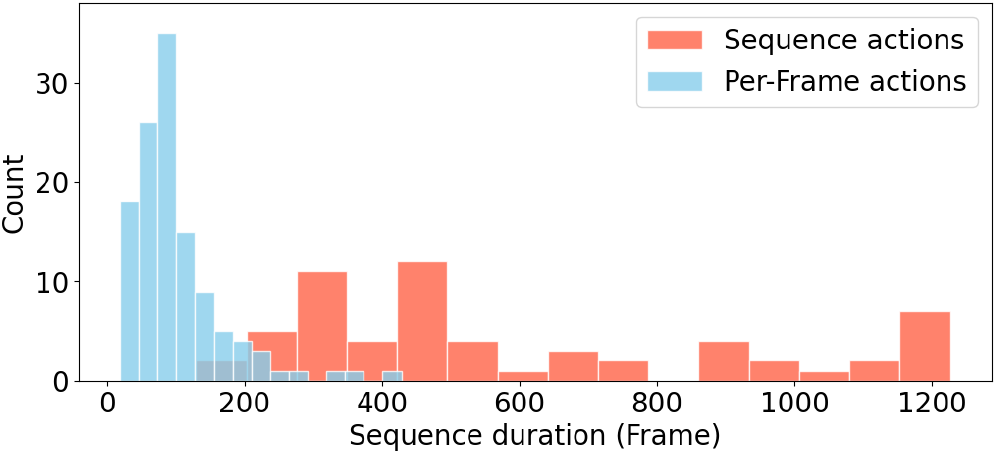
Distribution of Sequence-Level and Frame-Level behaviors in hBABEL.

**Fig. S7.**
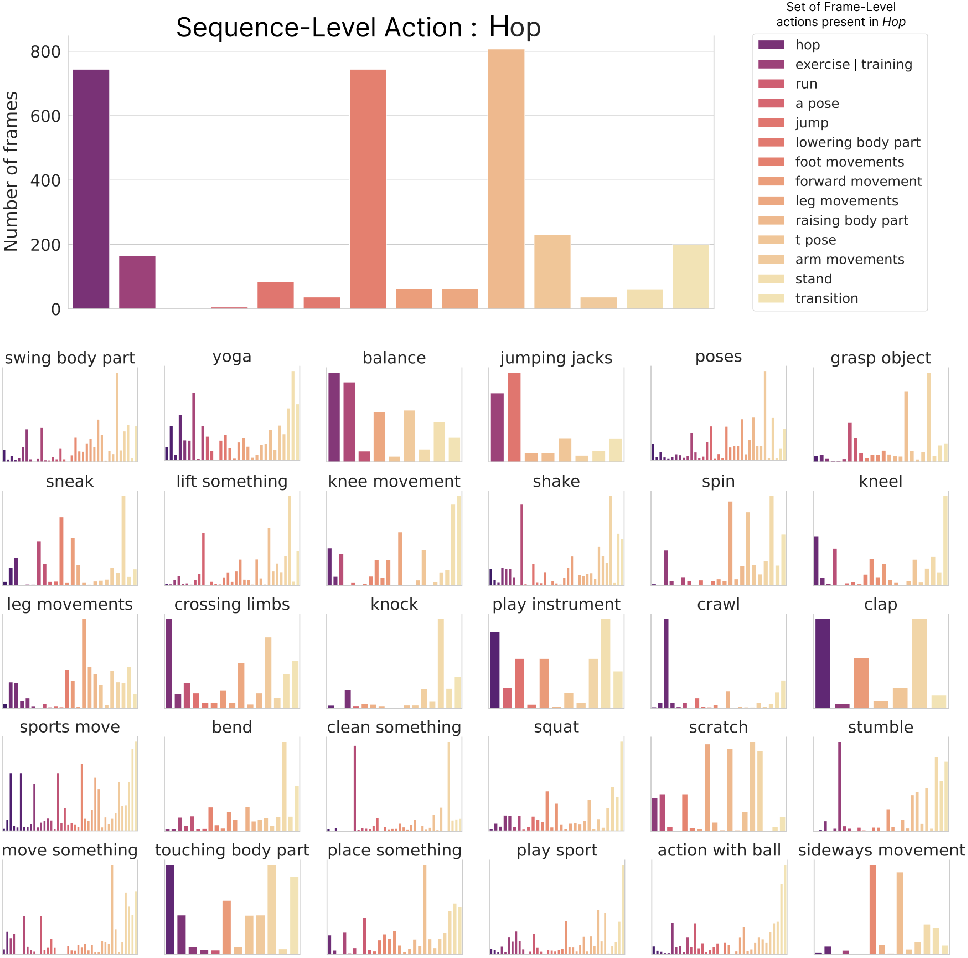
Distribution of Frame-Level actions per Sequence-Level actions. We show a selection of 31 Sequence-Level actions. Each plot represents the number of frames for each Frame-Level action per Sequence-Level action. Each color corresponds to a different Frame-Level action.

hBABEL is very challenging (no model reaches even 50% F1). This will hopefully inspire many algorithmic developments. It would also be great to annotate movemes, actions and activities consistent with Shot7M2 in the future.

### Supplementary Note B: h/BehaveMAE Implementation Details

In this section, we outline the specific design choices and hyperparameter settings used for the top-performing h/BehaveMAE models discussed in the main text. Additionally, we provide details on the baseline training setups for our hierarchical action segmentation benchmarks, Shot7M2 and hBABEL.

#### B.1. h/BehaveMAE architecture and pre-training

To construct a hierarchically interpretable model, careful consideration was given to the design of the hierarchical strides, which define how information is fused across different levels within the hBehaveMAE model (Table S2). Across all three datasets, we observed robust performance with 3-block architectures, incorporating local attention in the initial block. For the Shot7M2 dataset, we devised a spatio-temporal hierarchical model, beginning with small input token sizes (2×1×3 - representing a single keypoint over 2 frames) and gradually merging it into body parts, consisting of four spatially close keypoints, and then full poses with an increasing temporal window (resulting in a final token size of 8×1×72). Conversely, in the case of hBABEL, where the dataset’s labels are better characterized by their duration, we found that a temporally hierarchical model demonstrated the highest interpretability. For the MABe22 dataset, we started with full individual poses and progressively merged them into a larger temporal window and into the full group of animals, separately.

Overall, this also highlights the flexibility of the h/BehaveMAE framework for incorporating different design choices based on different datasets.

To ensure a fair comparison between hBehaveMAE and the non-hierarchical BehaveMAE, we maintained consistent architectural and training settings across both models. However, owing to its non-hierarchical nature, BehaveMAE requires fixing a specific scale, determined by the input token size. We conducted experiments with BehaveMAE models using various input scales, matching those of the corresponding hBehaveMAE model at each block. In the main text, we presented the best-performing BehaveMAE model overall, which utilized an input token size of 4×1×12 (corresponding to the mid-level scale of hBehaveMAE) for the Shot7M2 dataset (and a masking ratio of 80%). For hBABEL, we showcased the two most effective models, illustrating the challenge faced by single-scale models in achieving high performance across hierarchical tasks — where they excel either at frame-level or sequence-level tasks, but not both. In contrast, the hBehaveMAE model demonstrates an adept balance in performance across tasks.

The pre-training settings for both hBehaveMAE and BehaveMAE models can be found in Table S3.

#### B.2. Benchmarking on Shot7M2, hBABEL and MABe22

In order to build the pre-training dataset, we use a sliding window of 17 (for Shot7M2/MABe22) and 11 frames (for hBABEL), respectively, to create subsequences from the original sequences. We found that shorter integration windows, although leading to more training samples, contribute to overfitting of h/BehaveMAE, and hence lower performance on the downstream tasks. These subsquences, that are used as training samples, are defined by an input length. In case of shorter lengths, they are padded equally at the beginning and the end of the sequences with copies of the first/last frame.

We experimented with with different ways of extracting h/BehaveMAE’s final representations used for the linear probing in the downstream tasks (Shot7M2 and MABe22):

a. As the benchmarks we evaluate BehaveMAE on require one embedding per time-step, we extract the representation at time *t* by centering the frame in the sequence of length *l* seen during training, to include equal temporal context from the past and the future.
b. To increase robustness, we padded the beginning and the end of every sequence with *l*{2 and created subsequences of length *l* with a sliding window of 1. These are fed into the BehaveMAE model giving *l* embeddings for every time step *t* which are then averaged to create the final latent.

We found that the second method gave better results and is used in all results unless otherwise noted. To parallelize the evaluation of embeddings, we used GNU parallel (99).

#### B.3. Baselines on Shot7M2/hBABEL Training settings for PCA and TempPCA

On Shot7M2, we fit a PCA model using 64 components. The only difference between PCA and TempPCA is the dimensionality of the data in the input, in PCA the input data corresponds to all keypoints in 3D for one given frame whereas TempPCA takes as input subsequences (as described in section B.2) each having dimension 26 × 3 × *l*. We had better performances when choosing a sliding window of *l* = 5 frames. After fitting, we reach 99.9% of explained variance with PCA and 98.4% of explained variance with TempPCA.

**Fig. S8.**
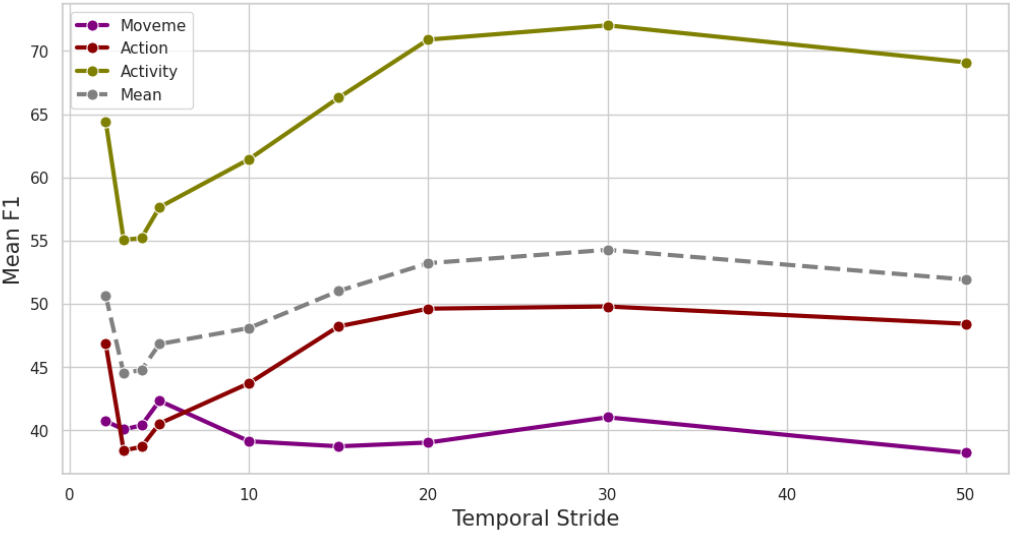
Performance vs. temporal stride for TVAE. Temporal stride is given as the number of input frames to the TVAE model.

##### Training settings for TVAE

We trained a TVAE model on Shot7M2 and hBABEL using the model and hyper-parameters implemented as baseline in the the MABe22 challenge (28). We first applied a SVD with 7 components to the centered and normalized data. The training process consisted of 300 epochs with a learning rate of 1*e* ^−5^ for Shot7M2 and 1*e* ^−4^ for hBABEL. A temporal stride of 30 frames centered around the represented frame was used. We optimized the learning rate and temporal stride through a grid search approach, varying the learning rates from 1*e* ^−3^ to 1*e* ^−5^ and temporal strides from 1 frame to 50 frames. We found an optimized temporal stride of 30 frames. Notably, we observed no significant improvement in movemes performance with a smaller temporal stride, despite a decrease in Activity and Action F1 scores (Figure S8).

**Table S2.** Architecture configurations for top-performing hBehaveMAE models. Input token size and hierarchical strides specify how the hierarchy is built up. #Dims, #Blocks and #Heads show the hidden dimension size, the number of hierarchical encoder blocks and heads in each block, respectively.

| Benchmark | Shot7M2 | hBABEL | MABe22 |
| --- | --- | --- | --- |
| Input token size | 2x1x3 | 3x1x75 | 3x1x24 |
| Hierarchical strides | [2x1x4 - 2x1x6] | [5x1x1 - 3x1x1] | [5x1x1 - 1x3x1] |
| Input sequence length | 400 | 450 | 900 |
| #Dims | [96 - 192 - 384] | [128 - 256 - 512] | [128 - 256 - 512] |
| #Blocks | [2 - 3 - 4] | [2 - 3 - 4] | [3 - 4 - 5] |
| #Heads | [2 - 4 - 8] | [2 - 4 - 8] | [2 - 4 - 8] |
| Attention | [Local - Global - Global] | [Local - Global - Global] | [Local - Global - Global] |
| Masking ratio | 70% | 75% | 87.5% |
| Mask token size | 8x1x72 | 45x1x75 | 15x3x24 |
| #Params | 8.8M | 15.4M | 20.4M |

**Table S3.** Pre-training settings for h/BehaveMAE models. reported in the main text.

| Configuration | Shot7M2 | hBABEL | MABe22 |
| --- | --- | --- | --- |
| Optimizer | AdamW (92) | AdamW (92) | AdamW (92) |
| Optimizer momentum | $\beta_1, \beta_2 = 0.9, 0.95$ (96) | $\beta_1, \beta_2 = 0.9, 0.95$ (96) | $\beta_1, \beta_2 = 0.9, 0.95$ (96) |
| Weight decay | 0.05 | 0.05 | 0.05 |
| Learning rate | 1.6e-4 | 1.6e-4 | 1.6e-4 |
| Learning rate schedule | cosine decay (97) | cosine decay (97) | cosine decay (97) |
| Warmup epochs (98) | 40 | 40 | 40 |
| Gradient clipping | 0.02 | 0.02 | 0.02 |
| Training epochs | 200 | 200 | 200 |
| Batch size | 512 | 512 | 1536 |
| Augmentation | - | - | Noise, Mirror., Rot. (93) |

##### Training settings for TS2Vec

TS2Vec (62) utilizes two types of contrastive losses for representation learning. The first loss contrasts a sequence with all others in a batch, treating subsequences from the same sequence as positives (instance contrastive). The second operates within a single time series, with nearby time points as positives and the rest as negatives (temporal contrastive). To find the best settings, we experimented with the different loss variants, resulting in models TS2Vec-I, TS2Vec-T, and TS2Vec-IT, following the baseline training in Azabou *et al*. (61).

On Shot7M2 we obtained the best model with both losses and a small number of training epochs (Table S4).

On hBABEL, using the instance contrastive loss alone works better, while a subsequence length of 300 during training showed higher performance than the sequence length used for h/BehaveMAE (450) (Table S5). In the main text we report the best results.

**Table S4.** Experiments for TS2Vec on Shot7M2. All models are trained with full sequences as input.

| Parameters |  |  | Results |  |  |  |
| --- | --- | --- | --- | --- | --- | --- |
| model | #epochs | hidden dim | Mean F1 | Moveme F1 | Action F1 | Activity F1 |
| TS2Vec-I | 50 | 512 | 59.1 | 59.8 | 49.4 | 68.1 |
| TS2Vec-I | 100 | 512 | 56.9 | 57.6 | 47.7 | 65.3 |
| TS2Vec-IT | 200 | 128 | 54.0 | 55.0 | 45.5 | 61.6 |
| TS2Vec-IT | 200 | 256 | 58.9 | 59.5 | 49.4 | 67.8 |
| TS2Vec-IT | 30 | 512 | **59.6** | **60.2** | **50.0** | **68.6** |
| TS2Vec-IT | 50 | 512 | 59.3 | 59.8 | 49.9 | 68.1 |
| TS2Vec-IT | 100 | 512 | 59.3 | 59.8 | 49.8 | 68.4 |

**Table S5.** Experiments for TS2Vec on hBABEL. All models are trained with a hidden dimension of 512 and for 100 epochs.

| Parameters |  | Results |  |
| --- | --- | --- | --- |
| model | input seq len | Frame-level | Sequence-level |
| TS2Vec-T | 300 | 16.5 | 10.3 |
| TS2Vec-I | 300 | **17.3** | **10.8** |
| TS2Vec-I | 450 | 16.3 | 10.6 |
| TS2Vec-IT | 300 | 16.3 | 10.2 |
| TS2Vec-IT | 450 | 16.4 | 10.0 |

##### Training settings for BAMS

We trained BAMS (61) models using the official implementation adapted for 1 individual on Shot7M2 and hBABEL. BAMS possesses a short-term and a long-term component. We optimized different short-term and long-term dilation coefficients (*st* and *lt*), as well as different model sizes, with the default model (large) having the following number of channels [64,64,32,32] and [64,64,64,32,32] for the short term and the long term encoder respectively, while the smaller model (small) having [64,32,32] and [64,64,32,32], respectively. With the small model, a short-term dilation of 2 corresponds to a receptive field size of 29, while a long-term dilation of 3 results in a receptive field of size 161. Each model on Shot7M2 (Table S6) was trained with a learning rate of 1*e* ^− 4^ and a sequence length of 1800 (full sequence). On hBABEL, we differentiate embeddings taken from the short-term encoder, the long-term encoder or the concatenation of both. In order to cope with different episode durations, we experimented with different sets of sequence lengths fed into the model during training. We note that we ran into out-of-memory errors when using the default settings (from GitHub) with very low batch sizes, thus limiting the amount of experimentation. Overall, training is relatively slow (2 days per model), which also limited our hyperparameter optimization.

**Table S6.** Experiments for BAMS on Shot7M2. F1-scores for BAMS models with different long-term (lt) and short-term (st) dilation coefficients.

| Parameters |  |  | Results |  |  |  |
| --- | --- | --- | --- | --- | --- | --- |
| *lt* | *st* | *Model size* | *Mean F1* | *Moveme F1* | *Action F1* | *Activity F1* |
| 2 | 1 | large | 51.3 | 48.8 | 41.8 | 63.2 |
| 3 | 2 | small | **53.7** | **48.6** | **45.3** | **67.1** |
| 4 | 2 | small | 47.9 | 40.5 | 39.1 | 64.0 |
| 4 | 2 | large | 52.9 | 47.9 | 44.0 | 66.7 |

**Table S7.** Experiments for BAMS on hBABEL. BAMS models with input sequence length of 1000 frames during tranining worked best, while we did not observe hierarchical interpretability. The combined embeddings always outperform both short-term and long-term embeddings.

| Parameters |  | Results |  |
| --- | --- | --- | --- |
| *Input seq len* | *Level* | *Frame-level* | *Sequence-level* |
| 300 | short-term | 8.1 | 7.3 |
| 300 | long-term | 10.0 | 9.0 |
| 300 | combined | 10.2 | 9.2 |
| 450 | short-term | 8.3 | 7.4 |
| 450 | long-term | 9.5 | 8.7 |
| 450 | combined | 9.7 | 8.8 |
| 1000 | short-term | 8.6 | 7.8 |
| 1000 | long-term | 10.4 | 9.1 |
| 1000 | combined | **10.8** | **9.4** |

### Supplementary Note C: Detailed Results

#### C.1. Detailed results on MABe22

Here, we present detailed results of h/BehaveMAE’s performance on the MABe22 representation learning benchmark (28) (Table S8). While our models outperform previous SotA methods on frame-level tasks, they also demonstrate competitive or improved performance on sequence-level tasks. Notably, we do not display results for oral-ear, oral-genital, and oral-contact scores, as these subtasks remain unsolved by existing methods. These results reflect the practical utility of our approach in capturing complex behavioral patterns directly from raw pose data without the need for extensive feature engineering.

#### C.2. Detailed results on Shot7M2

We additionally show detailed results for all action classes in the Shot7M2 dataset. hBehaveMAE achieves the highest performance across many movemes, nearly all actions and all activities (Table S9). Overall, hBehaveMAE has the best averaged F1-score for activities with 80.9%, actions with 58.5%, and movemes with 67.3%. We observe that the classification of movemes deteriorates with higher-level embeddings, while the opposite effect is seen with activities, showing that hBehaveMAE is able to effectively learn a hierarchical structure of action categories. We note that some classes such as *Spin* or *Switch* are hardly decoded for all models due to their rare frequency. In addition, hBehaveMAE balances the performance across the categories well, a characteristic not observed in other models, such as the non-hierarchical BehaveMAE (−6.3% points on movemes) and PCA-30 (−7.7% points on actions, and −12.6% points on activities).

### Supplementary Note D: Unsupervised analysis of hBehaveMAE Latents

We investigate the hBehaveMAE embeddings without using labels for assessing linear projections of the latent spaces (linear probing). We clustered hBehaveMAE latents on Shot7M2 with k-means and assigned each cluster a class label based on the most frequently appearing class within the cluster. Using this assignment, we sorted the clusters for both low-level and high-level embeddings. We found that clusters could often be assigned to single behavior labels or sensible combinations thereof. Consistent with our linear probing results, we find that more clusters for high-level embeddings represent activities vs. low-level embeddings (28 vs 9), while low-level ones better cover movemes (55 vs 35), suggesting that hBehaveMAE learns a hierarchical representation of behavior. (Figure S9)

### Supplementary Note E: Ablations

All the following analysis are performed by ablating the base model that uses multi-scale decoding, a masking ratio of 80% and local attention in its first block. The ablations were carried out on Shot7M2. Colors in tables correspond to the origin of the embeddings in the hierarchical levels of hBehaveMAE.

#### E.1. Reconstruction target

We first investigated the impact of the target scale on performance. Specifically, we chose to reconstruct poses at the 1-st scale, corresponding to the scale of the input tokens, which have a temporal stride of 2 in the base model. Reconstructing all frames from the input sequence yield no discernible performance increase and introduce unnecessary overhead. Conversely, reconstructing poses at the highest scale led to a drop in performance, attributed to the diminished resolution at this level (Table S10).

**Table S8.** Linear readouts of 10 subtasks of MABe22. BehaveMAE models reach state-of-the art performance, exceeding especially in important frame-level tasks.: marks the inclusion of the test data during pre-training.

| Method | Feat. Eng. | Sequence-level subtasks |  |  |  |
| --- | --- | --- | --- | --- | --- |
| | | Day (↓) | Time (↓) | Strain | Lights |
| PCA |  | .09416 | .09445 | 51.60 | 54.65 |
| TVAE |  | .09403 | .09442 | 52.98 | 56.80 |
| TS2Vec |  | .09380 | .09422 | 57.12 | 65.60 |
| T-Perceiver | ✓ | .09322 | .09323 | 69.81 | 69.68 |
| T-GPT† |  | .09269 | .09384 | 64.45 | 65.39 |
| T-PointNet | ✓ | .09275 | .09320 | 66.01 | 67.15 |
| T-BERT | ✓ | .09262 | .09276 | 78.63 | 68.84 |
| BAMS† | ✓ | **.09094** | **.08989** | **88.23** | 72.00 |
| **BehaveMAE†** |  | **.09094** | <u>.09028</u> | <u>87.58</u> | **74.38** |
| **hBehaveMAE†** |  | <u>.09186</u> | .09176 | 87.14 | <u>73.43</u> |

| Method | Approach | Frame-level subtasks |  |  |  |  |
| --- | --- | --- | --- | --- | --- | --- |
|  |  | Chase | Close | Contact | Huddle | Watching |
| PCA | 0.86 | 0.14 | 49.27 | 37.87 | 12.71 | 6.65 |
| TVAE | 1.07 | 0.45 | 59.33 | 44.77 | 21.96 | 10.20 |
| TS2Vec | 1.29 | 0.66 | 59.53 | 46.13 | 24.74 | 12.37 |
| T-Perceiver | 1.57 | 1.27 | 60.84 | 47.81 | 28.32 | 16.42 |
| T-GPT† | 1.73 | 0.64 | 69.05 | 55.78 | 23.80 | 17.86 |
| T-PointNet | 2.56 | <u>4.57</u> | 70.68 | 55.96 | 21.23 | 15.61 |
| T-BERT | 1.80 | 0.87 | 70.22 | 55.84 | 30.24 | 17.27 |
| BAMS† | 2.74 | 1.89 | 67.22 | 53.43 | 31.43 | <u>18.15</u> |
| **BehaveMAE†** | **3.05** | **4.92** | <u>72.03</u> | <u>58.54</u> | <u>33.60</u> | 17.19 |
| **hBehaveMAE†** | <u>2.80</u> | 4.01 | **73.57** | **60.55** | **35.05** | **20.29** |

**Fig. S9.**
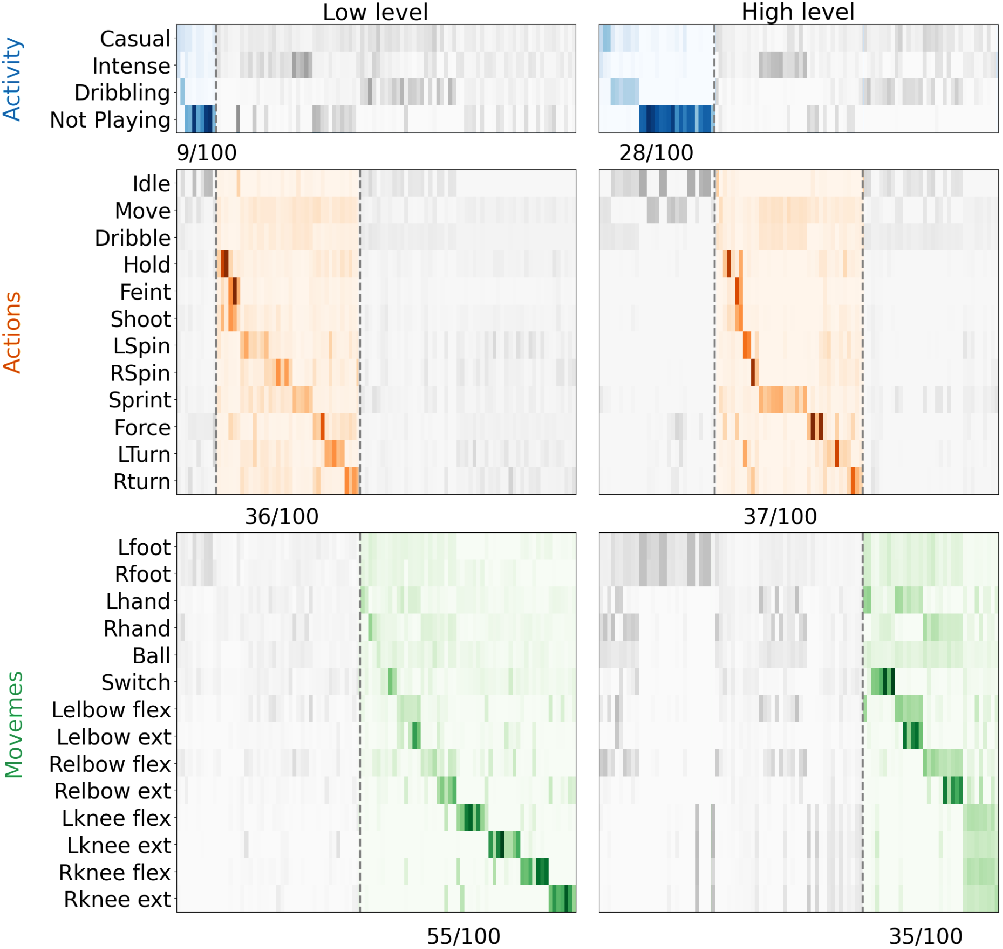
Visualization of the co-occurrences between clusters of hBehaveMAE latents and Shot7M2 class labels. Numbers correspond to fractions of assigned clusters to hierarchical category.

#### E.2. Drop path

Ablating the droppath rate reveals that dropping some paths during training results in marginal performance gains for actions and activities, albeit at the cost of decreased performance for movemes, mirroring the effects observed with higher masking rates (Table S11). Therefore, to maintain a balanced performance across all hierarchical categories, we kept no droppath regularization.

#### E.3. Model size

We furthermore varied both the encoder size (hidden dimensions and attention heads) and decoder size (depth and attention heads) of the base model (Table S12 and S13).

We observe that increasing the encoder size leads to a drop in overall performance, particularly in achieving balanced performance across all three categories. This imbalance is likely attributed to the increased variability in embedding sizes across categories. Considering the relatively modest size of our datasets, we hence keep a rather lightweight encoder to ensure computational efficiency. Additionally, we find that deeper decoders do not yield performance gains, possibly due to the evaluation through linear probing. As a result, we prioritize simplicity in the decoder architecture.

#### E.4. Loss function

hBehaveMAE is robust with regard to the choice of loss function, with mean squared error (MSE) showing best performance (Table S14).

**Table S9.**
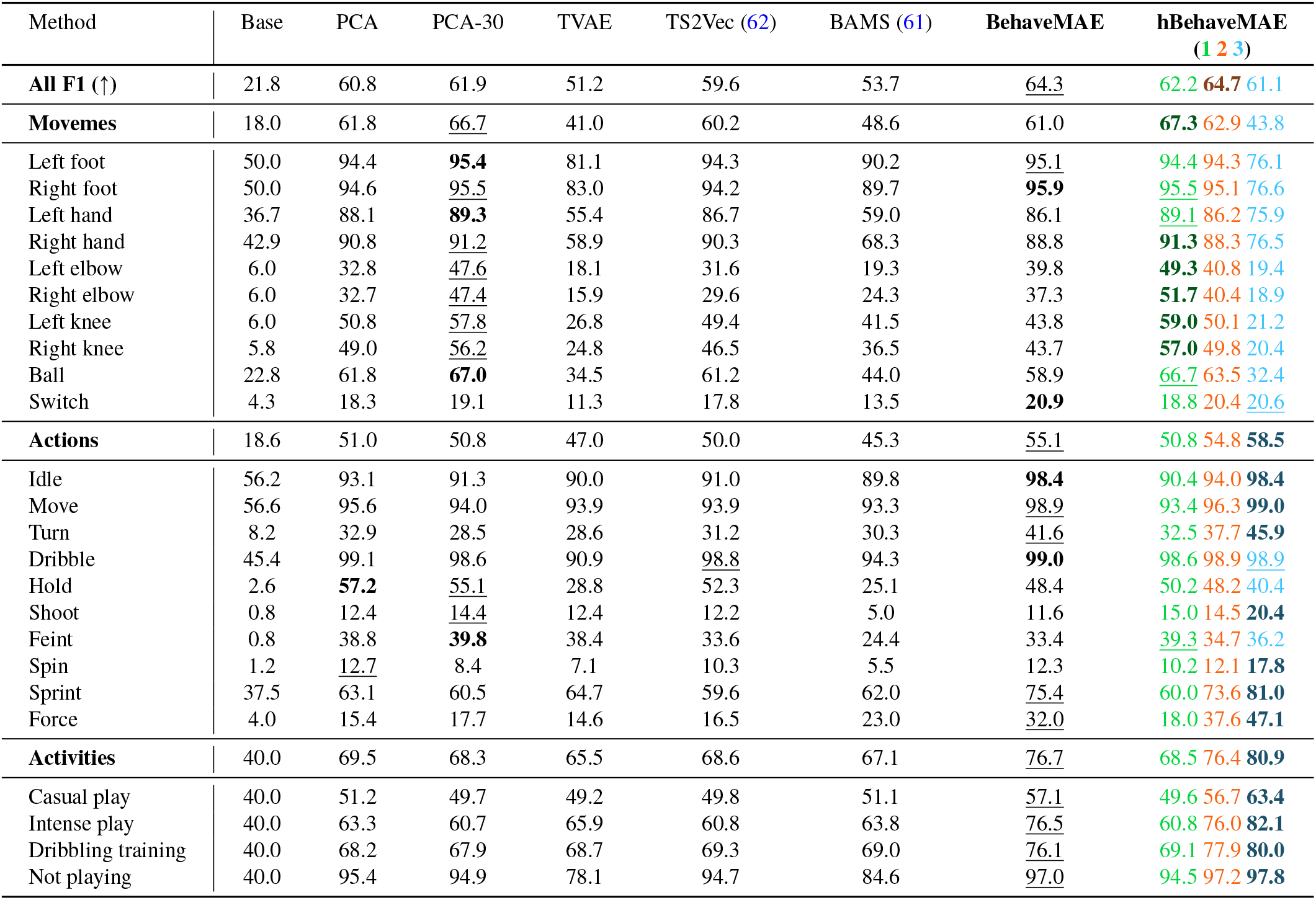
Results on Shot7M2: Our hierarchical BehaveMAE outperforms baseline methods across all three categories movemes, actions, and activities. We show class performance for every hierarchical level and hence reveal where an action class falls into the models hierarchy. The *All F1* score is calculated as the average over each behavioral scale average score. BehaveMAE scores are obtained from embeddings of layer 4 (its best).

| Method | Base | PCA | PCA-30 | TVAE | TS2Vec (62) | BAMS (61) | BehaveMAE | hBehaveMAE<br>(1 2 3) |
| --- | --- | --- | --- | --- | --- | --- | --- | --- |
| **All F1 (↑)** | 21.8 | 60.8 | 61.9 | 51.2 | 59.6 | 53.7 | <u>64.3</u> | <u>62.2</u> **64.7** 61.1 |
| **Movemes** | 18.0 | 61.8 | <u>66.7</u> | 41.0 | 60.2 | 48.6 | 61.0 | **67.3** 62.9 43.8 |
| Left foot | 50.0 | 94.4 | **95.4** | 81.1 | 94.3 | 90.2 | <u>95.1</u> | 94.4 94.3 76.1 |
| Right foot | 50.0 | 94.6 | <u>95.5</u> | 83.0 | 94.2 | 89.7 | **95.9** | <u>95.5</u> 95.1 76.6 |
| Left hand | 36.7 | 88.1 | **89.3** | 55.4 | 86.7 | 59.0 | 86.1 | <u>89.1</u> 86.2 75.9 |
| Right hand | 42.9 | 90.8 | <u>91.2</u> | 58.9 | 90.3 | 68.3 | 88.8 | **91.3** 88.3 76.5 |
| Left elbow | 6.0 | 32.8 | <u>47.6</u> | 18.1 | 31.6 | 19.3 | 39.8 | **49.3** 40.8 19.4 |
| Right elbow | 6.0 | 32.7 | <u>47.4</u> | 15.9 | 29.6 | 24.3 | 37.3 | **51.7** 40.4 18.9 |
| Left knee | 6.0 | 50.8 | <u>57.8</u> | 26.8 | 49.4 | 41.5 | 43.8 | **59.0** 50.1 21.2 |
| Right knee | 5.8 | 49.0 | <u>56.2</u> | 24.8 | 46.5 | 36.5 | 43.7 | **57.0** 49.8 20.4 |
| Ball | 22.8 | 61.8 | **67.0** | 34.5 | 61.2 | 44.0 | 58.9 | <u>66.7</u> 63.5 32.4 |
| Switch | 4.3 | 18.3 | 19.1 | 11.3 | 17.8 | 13.5 | **20.9** | 18.8 20.4 <u>20.6</u> |
| **Actions** | 18.6 | 51.0 | 50.8 | 47.0 | 50.0 | 45.3 | <u>55.1</u> | <u>50.8</u> 54.8 **58.5** |
| Idle | 56.2 | 93.1 | 91.3 | 90.0 | 91.0 | 89.8 | **98.4** | 90.4 94.0 **98.4** |
| Move | 56.6 | 95.6 | 94.0 | 93.9 | 93.9 | 93.3 | 98.9 | 93.4 96.3 **99.0** |
| Turn | 8.2 | 32.9 | 28.5 | 28.6 | 31.2 | 30.3 | <u>41.6</u> | 32.5 37.7 **45.9** |
| Dribble | 45.4 | 99.1 | 98.6 | 90.9 | <u>98.8</u> | 94.3 | **99.0** | 98.6 98.9 <u>98.9</u> |
| Hold | 2.6 | **57.2** | <u>55.1</u> | 28.8 | 52.3 | 25.1 | 48.4 | 50.2 48.2 40.4 |
| Shoot | 0.8 | 12.4 | <u>14.4</u> | 12.4 | 12.2 | 5.0 | 11.6 | 15.0 14.5 **20.4** |
| Feint | 0.8 | 38.8 | **39.8** | 38.4 | 33.6 | 24.4 | 33.4 | <u>39.3</u> 34.7 <u>36.2</u> |
| Spin | 1.2 | <u>12.7</u> | 8.4 | 7.1 | 10.3 | 5.5 | 12.3 | 10.2 12.1 **17.8** |
| Sprint | 37.5 | <u>63.1</u> | 60.5 | 64.7 | 59.6 | 62.0 | <u>75.4</u> | 60.0 73.6 **81.0** |
| Force | 4.0 | 15.4 | 17.7 | 14.6 | 16.5 | 23.0 | <u>32.0</u> | 18.0 37.6 **47.1** |
| **Activities** | 40.0 | 69.5 | 68.3 | 65.5 | 68.6 | 67.1 | <u>76.7</u> | <u>68.5</u> 76.4 **80.9** |
| Casual play | 40.0 | 51.2 | 49.7 | 49.2 | 49.8 | 51.1 | <u>57.1</u> | 49.6 56.7 **63.4** |
| Intense play | 40.0 | 63.3 | 60.7 | 65.9 | 60.8 | 63.8 | <u>76.5</u> | 60.8 76.0 **82.1** |
| Dribbling training | 40.0 | 68.2 | 67.9 | 68.7 | 69.3 | 69.0 | <u>76.1</u> | 69.1 77.9 **80.0** |
| Not playing | 40.0 | 95.4 | 94.9 | 78.1 | 94.7 | 84.6 | <u>97.0</u> | 94.5 97.2 **97.8** |

**Table S10.**
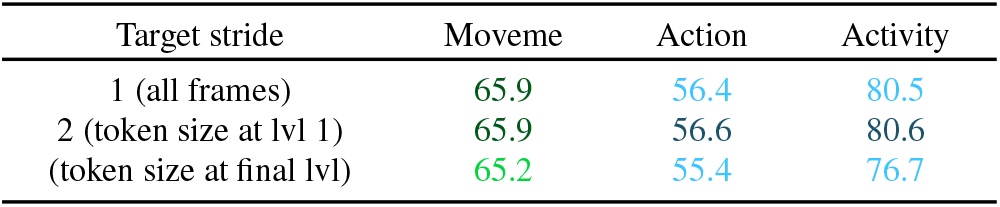
Temporal reconstruction target. Following the strategy of Ryali et al. (80), namely reconstructing the target at the 1-st scale works best also for hBehaveMAE.

| Target stride | Moveme | Action | Activity |
| --- | --- | --- | --- |
| 1 (all frames) | 65.9 | 56.4 | 80.5 |
| 2 (token size at lvl 1) | 65.9 | 56.6 | 80.6 |
| (token size at final lvl) | 65.2 | 55.4 | 76.7 |

**Table S11.**
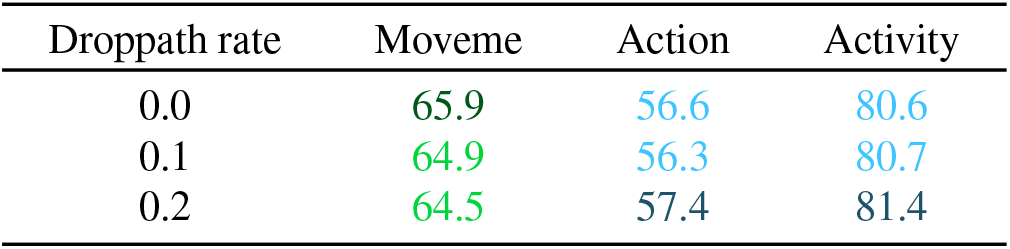
Drop path. Similar to higher masking rates, introducing drop path regularization helps for coarse-grained categories, but results in performance drops in movemes.

**Table S12.** Encoder size.

| Hidden dims | Att heads | # Params | Moveme | Action | Activity |
| --- | --- | --- | --- | --- | --- |
| 96-192-384 | 2-4-8 | 8.8M | 65.9 | 56.6 | 80.6 |
| 128-256-512 | 4-8-16 | 15.5M | 64.0 | 52.3 | 76.1 |
| 256-512-1024 | 4-8-16 | 60.9M | 56.0 | 52.1 | 83.4 |

**Table S13.** Decoder size.

| Depth | Att heads | Moveme | Action | Activity |
| --- | --- | --- | --- | --- |
| 1 | 1 | 65.9 | 56.6 | 80.6 |
| 2 | 2 | 65.6 | 54.8 | 80.0 |
| 4 | 4 | 66.3 | 52.3 | 78.8 |

**Table S14.** Loss function. We found that the standard MSE loss works best compared to other objective functions.

| Loss | Moveme | Action | Activity |
| --- | --- | --- | --- |
| Normalized MSE | 65.5 | 54.8 | 79.0 |
| MSE | 65.9 | 56.6 | 80.6 |
| Cosine Error | 65.7 | 54.8 | 79.0 |
| Smooth L1 | 66.0 | 55.9 | 80.3 |

#### E.5. Embedding size

We systematically varied the output embedding size of our base model from 16 to 128. Across all categories (movemes, actions, activities), we observed a consistent trend of increased performance when the embedding size is increased (Figure S10), as expected. Despite the improved performance with larger embedding sizes, we opted to set the maximum allowed embedding size in the Shot7M2 benchmark to 64 for practical reasons, ensuring that models maintain compact representations while still achieving reasonable performance across all categories. This choice also allows a meaningful comparison with PCA, which operates on input data with a dimensionality of 78.

**Fig. S10.**
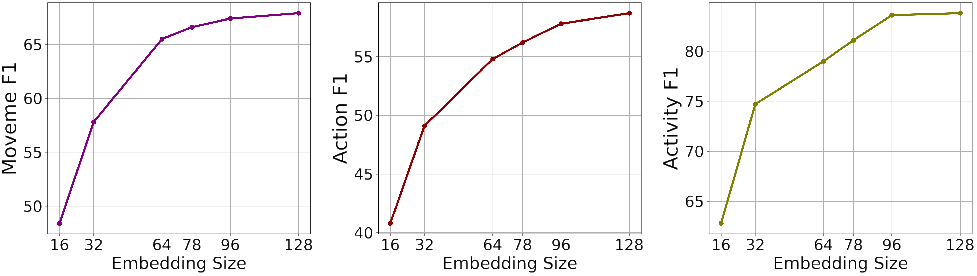
Embedding size. Increasing the embedding size results in improved performance across all categories on the Shot7M2 benchmark.

